# Bitbow: a digital format of Brainbow enables highly efficient neuronal lineage tracing and morphology reconstruction in single brains

**DOI:** 10.1101/2020.04.07.030593

**Authors:** Ye Li, Logan A Walker, Yimeng Zhao, Erica M Edwards, Nigel S Michki, Hon Pong Jimmy Cheng, Marya Ghazzi, Tiffany Y Chen, Maggie Chen, Douglas H Roossien, Dawen Cai

**Affiliations:** Department of Cell and Developmental Biology, University of Michigan Medical School; Biophysics, University of Michigan School of Literature, Science, and the Arts; Neuroscience Graduate Program, University of Michigan Medical School

## Abstract

Identifying the cellular origins and mapping the dendritic and axonal arbors of neurons have been century old quests to understand the heterogeneity among these brain cells. Classical chemical and genetic methods take advantage of light microscopy and sparse labeling to unambiguously, albeit inefficiently, trace a few neuronal lineages or reconstruct their morphologies in each sampled brain. To improve the analysis throughput, we designed Bitbow, a digital format of Brainbow which exponentially expands the color palette to provide tens of thousands of spectrally resolved unique labels. We generated transgenic Bitbow *Drosophila* lines, established statistical tools, and streamlined sample preparation, image processing and data analysis pipelines to allow conveniently mapping neural lineages, studying neuronal morphology and revealing neural network patterns with an unprecedented speed, scale and resolution.

## Introduction

Bilaterian nervous systems are built upon heterogeneous populations of neurons that form interconnected circuits. To understand the molecular and cellular mechanisms that lead to proper circuit formation, it is critical to elucidate the lineage origin and morphology formation of neurons. This is because lineages mark the outcome of neurogenesis, while morphology dictates the circuit structure by defining physical boundaries of the receptive and projective fields. Tremendous efforts have been made in the past century to take on these two fundamental quests in neuroscience, evolving from methodologies that can cope with one or a few neurons at a time, such as stochastic silver staining (Golgi’s method)^1,2^ and mosaic genetic labeling^3,4^, to multispectral labeling technologies (Brainbow) that can differentiate large population of neurons in the same tissue^5^.

Brainbow and Brainbow-like tools label neurons in distinct colors by expressing random ratios of different fluorophores, such as fluorescent proteins (FPs), upon genome recombination^6–8^. Reagents, including mice^5,9,10^, fruit flies^11–17^, zebrafish^18–21^, bacteria^22^, and viruses^10,23–25^ are now broadly available for lineage and morphology studies. In lineage studies, unique colors generated in the progenitor cells and inherited by their progenies were used to depict the clonal expansion process of adjacent lineages^9,15,19,26,27^. In morphology studies, the unique colors of neurites aided in identification of parallel projection patterns^21,28^ and confirming presynaptic inputs from multiple neurons converging to a common target^29–32^. However, current designs are often limited to generating up to tens of reliably distinguishable colors in a transgenic animal. The small unique color pool results in a high probability of labeling neighboring cells with the same color, therefore constraining the labeling density for neuronal morphology reconstructions. This makes it even more challenging to interpret lineage tracing results due to the need for unique colors to specify cells in the same lineage. In addition, distinguishing color variants differing by intensity levels in spectral channels is not reliable for lineage tracing because FP expression level may vary among cells in the same lineage.

One way to generate more unique labels for lineage tracing is to localize the same FPs to different subcellular compartments. In strategies such as CLoNe and MAGIC, Brainbow cassettes targeted to cytoplasm, cell membrane, nucleus, and/or mitochondria were co-electroporated with transposase for genome integration, which allowed the differentiation of neighboring progenies in chick and mouse embryos with fewer color collisions^26,27^. However, the number of expression cassettes being integrated in each cell is random in these experiments, leading to uncertainty in each color’s appearance probability which complicates quantitative analysis. The Raeppli strategy solves this problem by generating a transgenic *Drosophila* which utilizes 4 FPs to create up to 4 x 4 = 16 membrane and nucleus color combinations^16^. In parallel, strategies such as TIE-DYE and MultiColor FlpOut (MCFO) attempt to generate more color combinations by stochastically removing the expression stops from each FP module ^15,28^. While inserting 3 different modules into 3 genomic loci allows generating up to 2^3^-1=7 unique labels, it is difficult to insert more modules to more genomic loci in a single transgenic animal.

Here we present Bitbow, a digital format of Brainbow to greatly expand the unique color pool from a single transgenic cassette. Unlike the original Brainbow, whose FP choices are exclusive in one cassette, Bitbow allows each FP to independently express in an ON or OFF state upon recombination. Color coding by each FP’s binary status is similar to the information coding by each bit in computer memory, thus leading to the name Bitbow. In a recent study, we implemented the Bitbow1 design to target 5 spectrally distinct FPs to the nucleus for lineage tracing^33^. Here, we present novel Bitbow1 flies which encode up to 32,767 unique “colors” (Bitbow codes) in a single transgenic animal. This allows reliable lineage tracing without complicated statistical tests^33^. To better enable morphology tracing, we generated Bitbow2, which couples Bitbow1 to a self-regulating recombination mechanism. This enables generating consistent neuronal labeling by a simple cross of a Bitbow2 fly to an enhancer-Gal4 driver fly without the need for heat-shock.

## Results

### Characterization of Bitbow design in the *Drosophila* brain

To permit independent recombination of each FP, we utilized a pair of inverted FRT sites flanking a reversely positioned FP. downstream of a 10xUAS sequence and upstream of a polyadenylation sequence (**Fig. 1a**). This default OFF state guarantees a non-fluorescent expression. Upon Flp recombination, the flanked FP spins between the inverted FRT sites, resulting in either an ON or OFF state of expression driven by Gal4. Such a design exponentially increases the color-coding capacity with increasing numbers of bits (FPs) in the same transgenic animal (**Fig. 1b**), however, requires a transient Flp activity to ensure the recombination choice is stabilized, similar to the original Brainrow2 design^5^. In order to guarantee independent recombination between each FP, we used incompatible flanking FRT sequences. Other than the three previously-used incompatible Frt sites^10^, FRT-F3, FRT-5T2, and FRT-545, we identified FRT-F13, FRT-F14, and FRT-F15 as additional incompatible sites in a screen (**Fig. S1**)^34^. As FRT-F15 has lower recombination efficiency (data not shown), we ended up with a 5-bit Bitbow1.0 design that consists the other 5 FRT sites to control the independent recombination choices of mAmetrine, mTFP1, mNeonGreen, mKusabira-Orange2 and tdKatushaka2, respectively^35–39^. These FPs were chosen for their brightness, photo-stability, antigenicity, and spectral separation (**Fig. S2**). Finally, we made the cell membrane-targeting Bitbow1.0 (mBitbow1.0) fly to better reveal whole neuron morphology.

**Figure 1.**
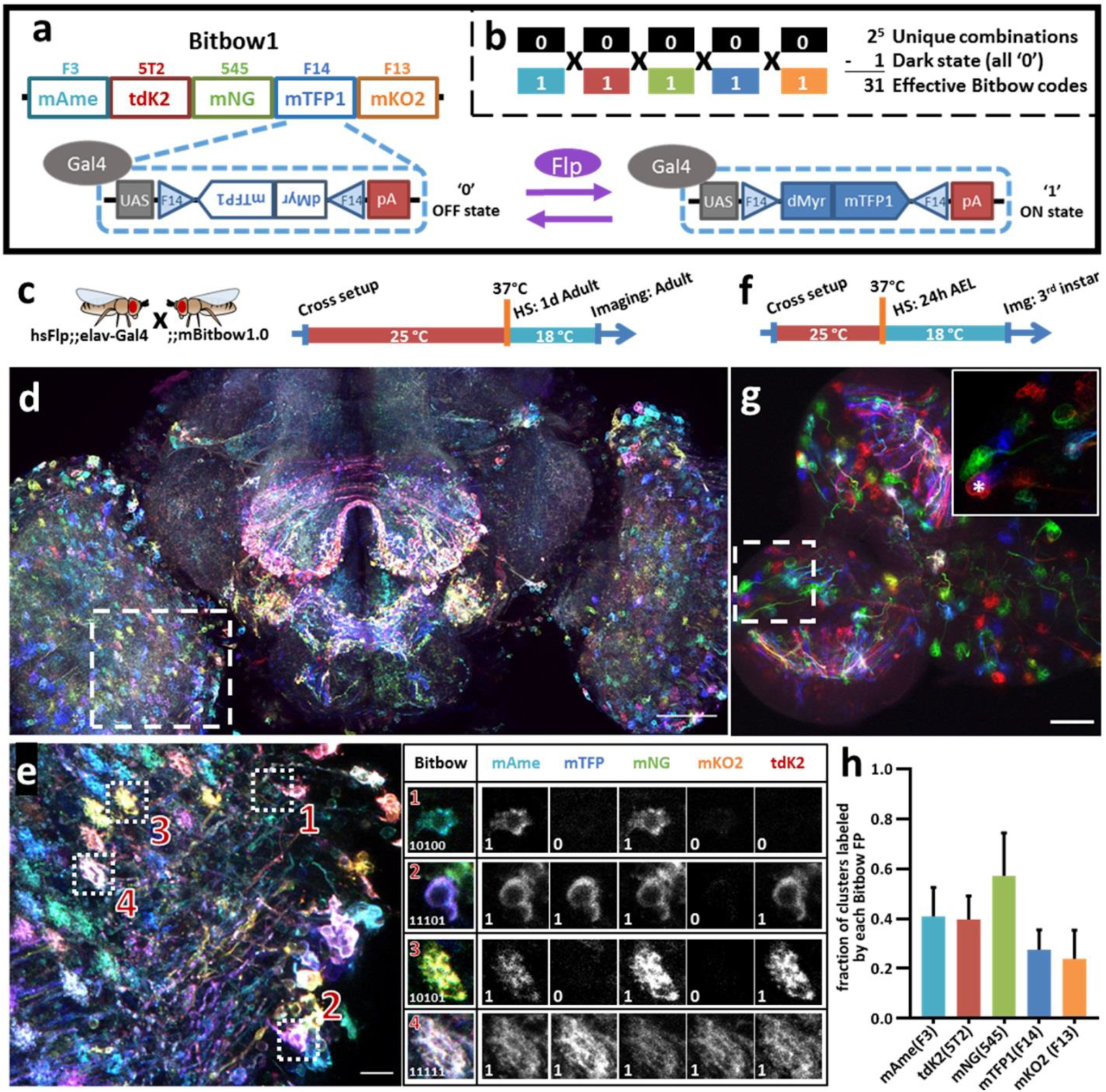
Bitbow1 design and characterization of labeling properties. **(a)** Schematic of Bitbow1 design. Five spectrally distinct fluorescent proteins (FPs) are separated by five pairs of reversely positioned orthogonal FRT sites. The mTFP/FRT-F14 module is shown in the dashed box. The FP’s open reading frame (ORF) is positioned in the reverse direction, corresponding to an default OFF state (‘0’). Upon Flp induced recombination, the FP’s ORF may spin to the forward direction for Gal4 driven expression, corresponding to an ON state (‘1’). **(b)** 31 Bitbow color codes could be generated in a single Bitbow1 brain. **(c)** A hsFlp;;elav-Gal4 driver fly was crossed to the mBitbow1.0 fly to examine the offspring expression in the nervous system upon heat-shock induced Flp activity. Experimental setups of adult heat shock-induced labeling. **(d)** Maximum intensity projection overview of an adult heat-chocked brain. **(e)** Left panel, enlarged boxed region in (**d**) showed individual neurons are labeled in distinct colors, i.e., Bitbow codes. Right panel, Bitbow codes of four selected optic lobe neurons’ somas or terminals. **(f)** Experimental setups of generating heat shock-induced Bitbow labeling in 3rd instar brains. **(g)** Maximum intensity projection overview of a 3rd instar larvae heat-chocked brain. Inset, the enlarged boxed region showed clusters of cells labeled in the same colors. Asterisk indicates a neuroblast. **(h)** Quantification of occurrence frequencies of each Bitbow color. Among all quantified clusters, the fraction of clusters containing each Bitbow color were displayed. 787 clusters from 6 brains are included. Error bars are SD. HS, heat-shock. Img, imaging. Scale bars: (**d**, **g**) 50μm, (**e**) 10μm.

Next, we crossed mBitbow1.0 flies to hsFlp;;elav-Gal4 driver flies to examine the offspring expression in the nervous system upon heat-shock induced transient Flp activity. When young adult offspring were heat-shocked at 1 day after eclosion and imaged at 3 days later (**Fig. 1c**), we observed individual neurons expressing unique combinations of Bitbow codes that can be denoted as a series of 5-bit 0/1 digits (**Fig. 1d**, **1e**). Increasing the number of heat-shocks (thus Flp activity) increased the total number of neurons being labeled and the number of FP species being expressed in each neuron (**Fig. S3**). Nonetheless, all 31 expected Bitbow codes were identified, albeit each of which was observed with a different frequency (**Fig. S4**). The appearance of strong and diverse Bitbow code labeling days after transient heat-shock also indicated that recombination outcomes induced by transient Flp activity were stable. Otherwise, all FPs would keep spinning so that they would all have some transcripts positioned in the forward direction to become fluorescent in all cells.

Depending on the timing of heat-shock, stochastic colors can be observed in neighboring neurons or clusters of neuronal progenies if recombination happens in postmitotic neurons or progenitor cells, respectively^26^. While post-eclosion heat-shock demonstrated the former situation, the later situation can be examined by heat-shocking at 24 hours after egg laid (24hr AEL, i.e., early 1st instar larval stage) and imaging at 72 hours post heat-shock, at the 3rd instar larval stage (**Fig. 1f**). Interestingly, while there are plenty of postmitotic neurons at the 1st instar larval stage, most neighboring neurons were labeled as cell clusters in the same Bitbow code (**Fig. 1g**). In addition, we always observed a much larger size neuroblast (NB, i.e., neural stem cell) being labeled in the same Bitbow code in each cluster (**Fig. 1g inset**, asterisk). Collectively, these observations suggested that under the heat-shock conditions optimized for larvae survival, recombination events mostly happened in the NBs and the recombination outcome did not change over time. Quantification of the expression frequency of each FP, i.e., the recombination rate of each FRT site, indicates that FRT-545 has the highest recombination rate, followed by FRT-F3, and FRT-5T2, while FRT-F14 and FRT-F13 have the similarly lowest among the five (**Fig. 1h**). This observation is not specific to the membrane targeting, but is consistent in other Bitbow1.0 flies (**Fig. S5** and detailed below).

### Targeting Bitbow FPs to multiple subcellular compartments permits high-throughput lineage tracing in the whole *Drosophila* brain without ambiguity

In a recent study, we specified the lineage relationships between pairs of *Drosophila* peripheral neurons using a nucleus-targeting Bitbow1.0 (nBitbow1.0) that can generate 31 unique Bitbow codes^33^. However, many more unique Bitbow codes are needed to unambiguously label the ^~^200 neuronal lineages in the *Drosophila* central brain. We made a membrane/nucleus double-targeted mnBitbow1.0 fly and an additional Golgi apparatus triple-targeted mngBitbow1.0 fly (**Fig. 2a**) to generate up to 1023 and 32,767 (**Fig. 2b**) unique Bitbow codes in the same brain, respectively.

**Figure 2.**
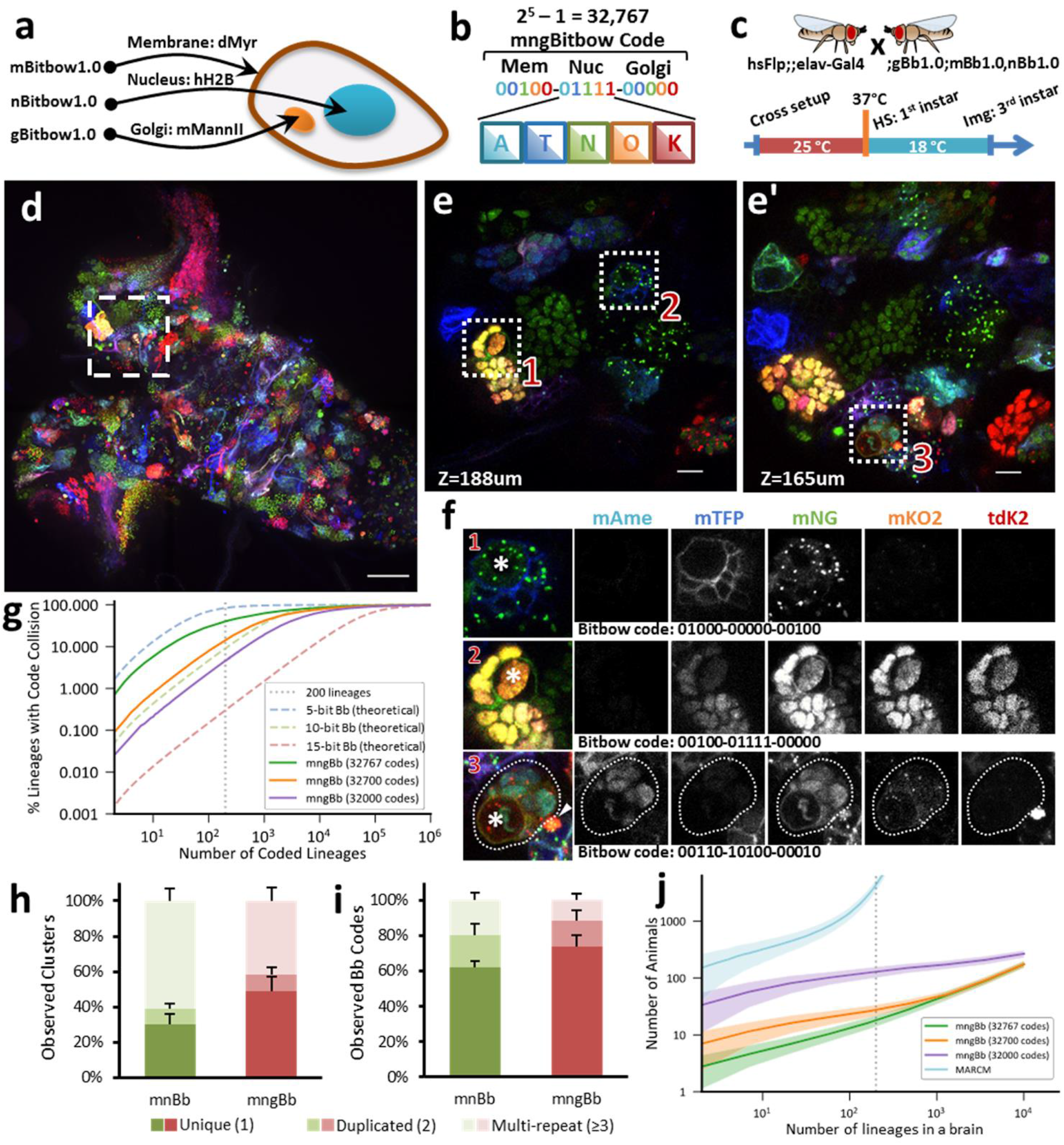
Targeting Bitbow fluorescent proteins (FPs) to multiple subcellular compartments enables high-throughput lineage tracing without ambiguity. **(a)** The same Bitbow FPs are targeted to cell membrane, nucleus or Golgi apparatus to generate spectrally-spatially resolvable Bitbow codes. **(b)** Up to 32,767 unique mngBitbow codes can be generated can be presented as 3 groups of 5-digit 0/1s in correspondence with the expression status of mAmetrine (A), mTFP1 (T), mNeonGreen (G), mKO2 (O) and tdKatushka2 (K). **(c)** Experimental setup of generating heat-shock induced mngBitbow1.0 labeling and imaging. **(d)** Maximum intensity projection of a 3rd instar mngBitbow1.0 brain. Scale bar, 50μm. **(e**, **e’)** Two confocal image slices corresponding to two different z positions in the boxed region in (**b**). Scale bar, 10μm. **(f)** 3 clusters marked in (**e**, **e’**) are assigned mngBitbow barcodes. Asterisks indicate the neuroblasts of each cluster. The arrowhead highlights an adjacent neuroblast labeled by a distinct mngBitbow code. **(g)** Simulation of Bitbow code collision in lineage mapping experiments. Dashed curve lines are simulations based on theoretical Bitbow code frequencies. Solid curve lines are simulations based on experimentally observed Bitbow code frequencies. Vertical dotted line corresponds to mapping all of the 200 lineages in a single adult *Drosophila* central brain. **(h)** Percentages of cell clusters that are uniquely labeled, or 2 of them, or >=3 of them are labeled by the same mnBitbow (286 clusters, 4 brains) or mngBitbow (577 clusters, 6 brains) codes in each brain. **(i)** Percentages of mnBitbow (N=80, 4 brains) or mngBitbow (N=240, 6 brains) codes that are expressed in 1, or 2, or >=3 clusters in each brain. Means and standard deviations (SD) are shown. **(j)** Monte Carlo simulations estimate the number of animals that are needed (y-axis) to sample all lineages at least once in animal brains that have given numbers of lineages (x-axis). Solid lines, means. Shaded lines, SD. HS, heat-shock. Img, imaging.

To examine the labeling efficacy, we crossed mnBitbow1.0 or mngBitbow1.0 flies to hsFlp;;elav-Gal4 flies, and performed the larval heat-shock experiment (**Fig. 2c**) to their offspring (**Fig. S5**, or **Fig. 2d**, respectively). As expected, we identified many cell clusters, in which all the cells were labeled by the same combinatorial Bitbow code (**Fig. 2d-2f**), which again indicated the transient Flp activity led to stable recombination outcome in the neural stem cells. Many of these Bitbow codes contain FPs in more than one subcellular compartments, which indicates that the repeated incompatible FRT sites inserted in distant chromosome locations are exempt from inter-Bitbow cassette recombination. In addition, these subcellular compartments are spatially well separated, even when the same FPs are expressed in different subcellular compartments in the same cell (**Fig. 2f**).

To estimate the theoretical ability to unambiguously distinguish Bitbow flies’ 200 lineages in the same *Drosophila* central brain, we ran a simulation to calculate how frequently the same Bitbow codes are seen in more than one lineage, i.e. the collision rate. The simulation shows that there would be a 84.5%, 9.1%, or 0.3% collision rate in a Bitbow fly that targets the 5 FPs to 1, 2, or 3 subcellular compartments, corresponding to 5-bit, 10-bit, or 15-bit Bitbow codes, respectively (**Fig. 2g** blue, green or red dashed lines, respectively). In other words, under uniformly random recombination conditions, we can identify any neuron’s lineage composition in the mngBitbow1.0 fly central brain with 99.7% confidence. To estimate the collision rate in real experiments, we conducted the early heat-shock experiment as shown in **Fig. 2c** with mnBitbow1.0 or mngBitbow1.0 flies. We plotted the percentages of cell clusters that are uniquely labeled, or 2 of them, or >=3 of them are labeled by the same Bitbow code in each brain. We found that the experimental collision rates of mnBitbow and mngBitbow fly brains are 69.8%±5.7% (mean±SD, 286 clusters from 4 brains) and 51.1%±8.5% (mean±SD, 577 clusters from 6 brains), respectively (**Fig. 2h**).

It seems desperate that the high collision rate would make even the mngBitbow1.0 fly useless for tracing neuronal lineages in the *Drosophila* central brain. However, we have shown that it is possible to develop a novel statistical method and apply it to the nBitbow1.0 flies to determine the lineage relationships between any two neighboring neurons in the *Drosophila* PNS^33^. Given that the mngBitbow1.0 fly generates much more unique Bitbow codes, we sought a different strategy to simplify the analysis yet ensure proper statistical power to unambiguously trace any neuronal lineage composition in the *Drosophila* central brain. We plotted the percentages of Bitbow codes that are expressed in 1, or 2, or >=3 clusters in each brain (**Fig. 2i**). We found that the majority of labeling collisions were contributed by a small number of Bitbow codes, most among which have mNeonGreen being turned on (**Fig. S6**). To estimate the effect of the FP turn-on bias to the apparent Bitbow code collision rates, we quantified the relative recombination frequencies of the incompatible FRT sites in mngBitbow1.0 (**Fig. S7a**), calculated the empirical frequencies of all 32,767 mngBitbow codes (**Fig. S7b**), and used the empirical frequencies to run the same simulation as shown above (**Fig. 2g**). We found that mngBitbow1.0’s experimental collision rate was estimated as 40.3% for 200 lineages (**Fig. 2g** green solid line), and a small number of codes appeared much more frequently, which contributed to most of the high collision events (**Fig. S7c**). Next, we excluded the most frequent 67 or 767 mngBitbow codes from the simulation and found the collision rate was decreased to 14.3% or 4.6%, respectively (**Fig. 2g**, orange, and purple solid lines). In other words, we have over 85.7% or 95.4% confidence to call any neurons belonging to the same lineage using the pool of 32,700 or 32,000 unique mngBitbow codes, respectively.

Encouraged by mngBitbow’s potential in determining lineage relationships with high confidence, we ran another simulation to estimate the number of animals needed to thoroughly survey the lineage relationship of any Gal4-driver labeled neurons across the whole central brain, i.e. every one of the 200 lineages needs to be observed at least once (**Fig. 2j**). We included the estimation for the popular method MARCM as a comparison^3^. In the simulation, we assumed a 48.08% lineage coverage for mngBitbow1.0 (577 clusters observed from six central brains containing an estimated total of 1200 neuronal lineages) and a 1% lineage coverage for MARCM to make sure no more than one lineage being labeled in each brain. This assumption underestimates the animal used in real MARCM experiments, that is because the same clonal patterns are normally required to be repeated more than once to confirm the labeling is indeed unique. Our simulation matches well with previous MARCM experiments ^40–42^, in which hundreds to thousands of brains were needed in one experiment (**Fig. 2j**, cyan line). Using mngBitbow1.0, only 28.3±6.4 flies (mean±SD) were needed to survey each of the 200 lineages at least once while achieving an overall >85% confidence in determining the lineage relationship between any neurons (**Fig. 2j**, orange line).

Next, we set out to use mngBitbow1.0 to determine the lineage relationships of serotonergic neurons in the *Drosophila* CNS. Serotonergic neurons are a group of ^~^100 neurons that have been shown to play critical roles in maintaining and regulating important neural functions in the fly, such as feeding, courtship, aggression, learning and memory^43–48^. Born in late embryonic stages, serotonergic neurons persist through larval, pupal and adult stages and form stereotypical clusters across the brain, especially in each hemi-segment of the ventral nerve cord (VNC, **Fig. 3a**)^49^. However, whether the serotonergic neurons in the same cluster originate from the same neural stem cell has not been experimentally determined. We crossed mngBitbow1.0 flies to hsFlp;TRH-Gal4; flies, heat-shocked the offspring embryos in an early neurogenesis period (0-4 hrs AEL), and imaged the serotonergic neurons in 3rd instar larva (**Fig. 3b**). We observed that all serotonergic neurons clustered in each hemi-segment of the VNC or in the central brain were labeled by distinct Bitbow codes (**Fig. 3c-3e**, **Fig. S8**), except on one occasion that two out of the four SE0 neurons near the esophagus hole were labeled by a Bitbow code of high collision rate (**Fig. S8**). We believe that the observation of adjacent serotonergic neurons being labeled in distinct Bitbow codes is not due to embryonic heat-shock, because in the control experiment utilizing a hsFlp;elav-Gal4; fly, many cell clusters labeled in the same Bitbow codes were observed (**Fig. S9**). In summary, our Bitbow study indicated that the majority of serotonergic neurons arise from distinct lineages.

**Figure 3.**
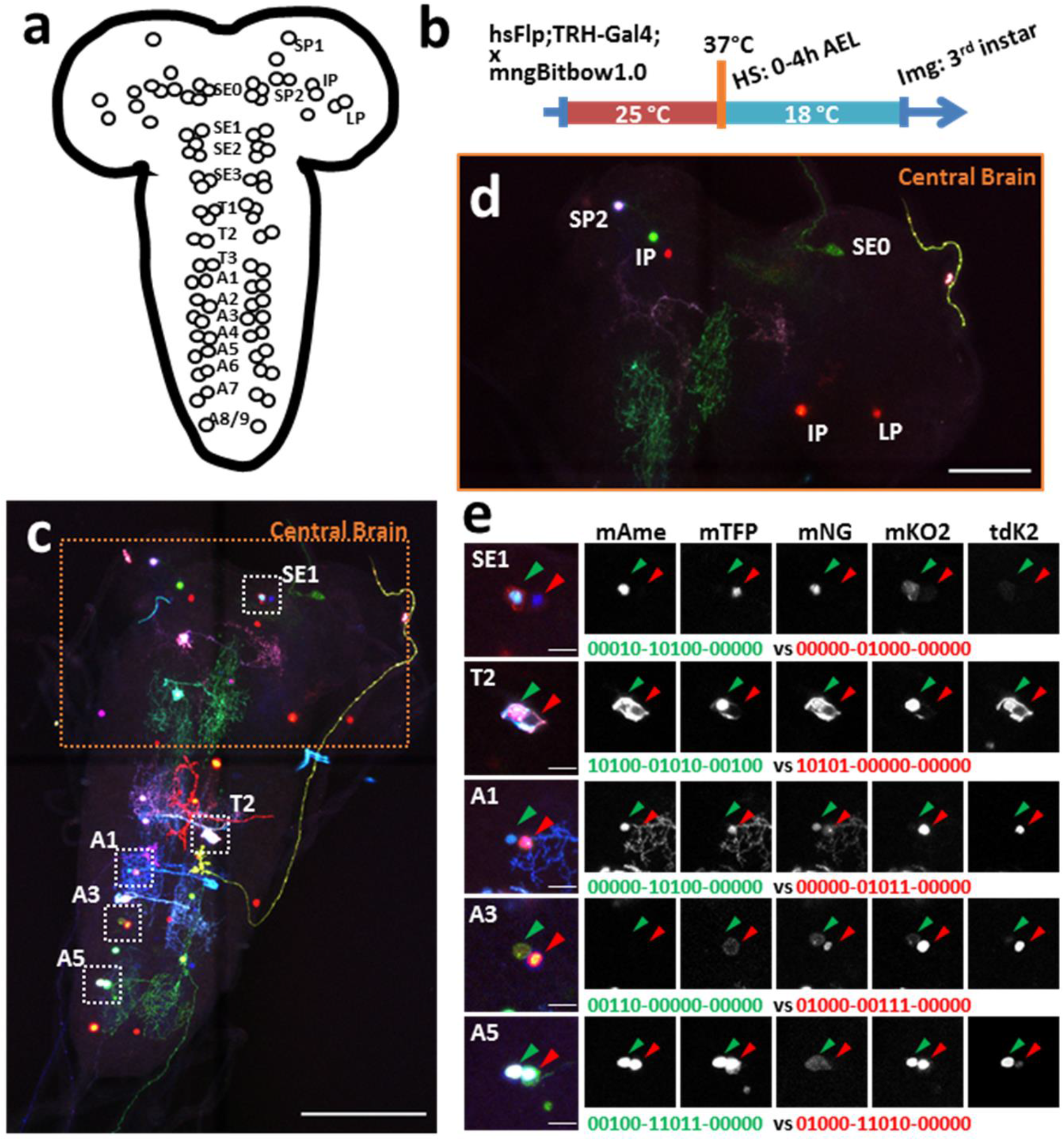
Mapping the lineage relationships between clustered serotonergic neurons by mngBitbow1.0. **(a)** Schematic of clustered serotonergic neuron somas (circles) in the 3rd instar brain. Neurons were named after their somas’ locations in the central brain regions or VNC segments. **(b)** Schematic of the experimental setup. **(c)** An example of mngBitbow1.0 labeled serotonergic neurons in a 3rd instar larval brain. **(d)** Enlarged view of the central brain area in (**c**). We found most of the serotonergic neurons were sparsely labeled in the central brain that did not form clusters. Cell cluster identities are estimated based on relative cell body location. **(e)** Enlarged boxed regions in (**c**) showed that pairs of serotonergic neurons in the same hemi-segments could be easily found. mngBitbow codes in green and red correspond to neurons marked by the green and red arrowheads, respectively. HS, heat-shock. Img, imaging. Scale bars: (**c**, **d**) 100μm, (**e**) 10μm.

### Bitbow2 enables broad neuron morphology labeling with a simple transgenic setup

While inducing Flp expression by heat-shock has the flexibility in controlling the timing of Bitbow1.0 recombination for lineage tracing, the relatively low Flp activity resulted in reduced color variation and labeling coverage, which constrains tracing morphology of postmitotic neurons (**Fig. S3**). Increasing heat-shock duration to increase Flp activity was not ideal, because the animals were challenged by stronger stress, which resulted in a lower survival rate (data not shown). In addition, the requirement of heat-shocks limited the use of Bitbow in combination with other temperature-dependent interrogations^50,51^. Finally, the hsFlp/enhancer-Gal4/Bitbow triple transgenes are more complicated to set up.

To overcome the above-mentioned limitations, we designed Bitbow2, in which a self-regulating Flp (srFlp) is added to Bitbow1 (**Fig. 4a**). The srFlp consists of a flippase cDNA flanked by a pair of FRT sites positioned in the same direction. Driven by the promoter of *Drosophila* neuronal Synaptobrevin (nSyb)^52^, this design permits a strong burst of neuronal-specific expression of flippase which recombines the FP modules to generate Bitbow codes and eventually excises out the flippase cDNA to prevent chromosome breaks caused by excessive recombination. To ensure sufficient amount of flippase being produced before its coding sequence being removed, we made mBitbow2.0 and mBitbow2.1, which utilized the less efficient FRT-F13 and FRT-F15 sites to lower the chance of self-excision, respectively.

**Figure 4.**
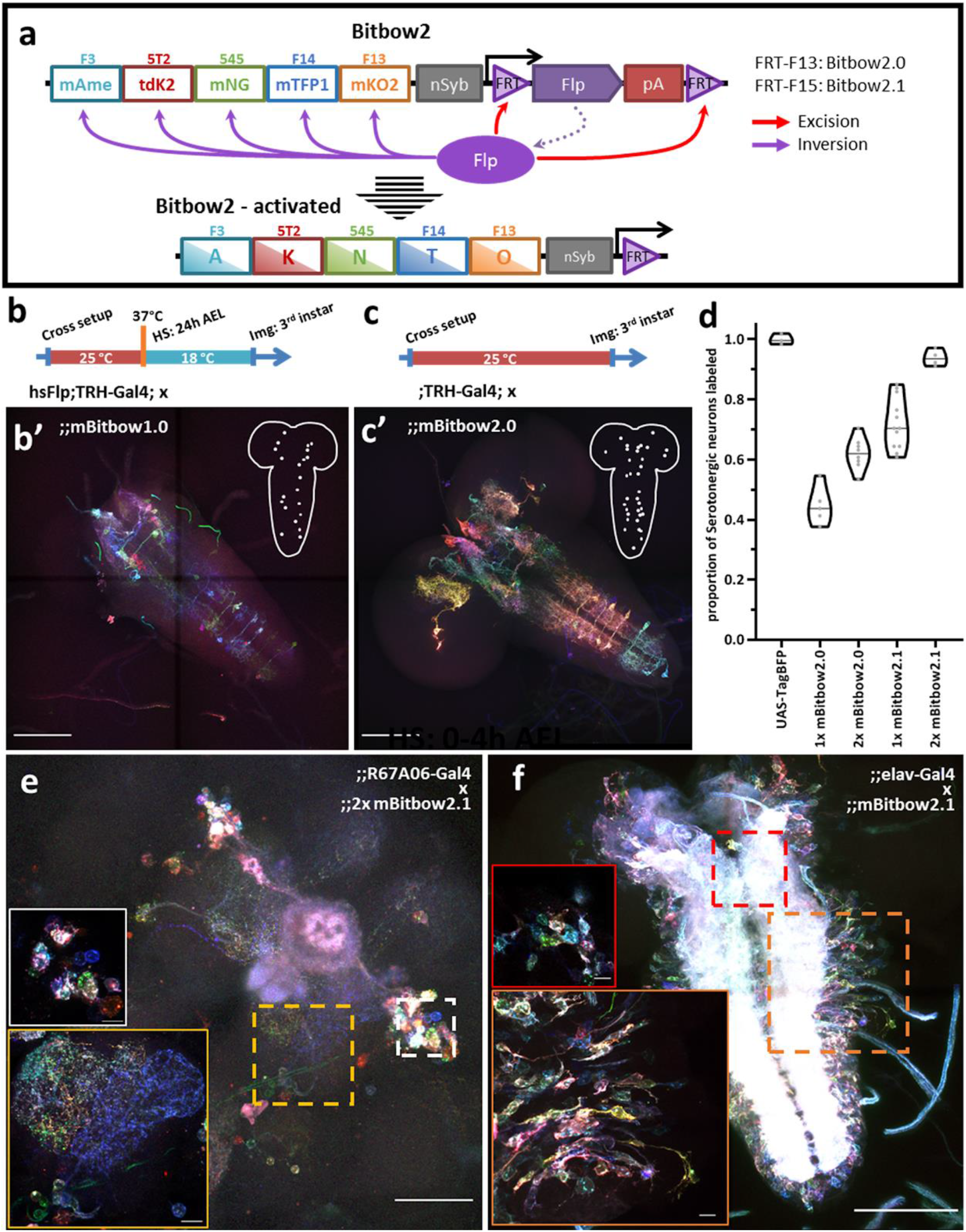
Bitbow2 enables broad neuron morphology labeling with a simple transgenic setup. **(a)** Schematic of Bitbow2 design. A self-regulating Flp module is added to ensure proper transient Flp activity without the need of an additional cross to the heat-shock Flp fly. Flp expression is driven by a neuron-specific n-Synaptobrevin (nSyb) promoter and terminated by self-excising between the flanking FRT sites, which have lower efficiency compared to those used in the Bitbow1 modules. This ensures proper Bitbow recombination before Flp self-excision to reach a stable genetic outcome. Compared to a **(b)** Bitbow1 labeling experiment, a **(c)** Bitbow2 labeling experiment requires only a direct cross to the TRH-Gal4 driver fly without the need of heat-shock. **(b’**, **c’)** indicate that mBitbow1.0 labeled fewer serotonergic neurons than Bitbow2.0 does. Inserted schematics indicate the somas of the labeled serotonergic neurons. **(d)** Quantification of the percentage of serotonergic neurons being labeled in different Bitbow2 flies, normalized to the labeling of a UAS-TagBFP fly. Each dot that overlays on the violin plots corresponds to the cell counting from one brain. **(e)** Adult neurons labeled in an offspring of the 2x mBitbow2.1 fly crossed to the R67A06-Gal4 fly. White and yellow insets show representative soma and neurite labeling, respectively. **(f)** Larva neurons labeled in an offspring of the 2x mBitbow2.1 fly crossed to the elav-Gal4 fly. Red and orange insets show neighboring neurons labeled in many distinct Bitbow colors in the central brain and in the VNC, respectively. HS, heat-shock. Img, imaging. Scale bars: (**b’**, **c’**, **f**) 100μm, (**e**) 50μm, (**e**, **f** inserts) 10μm.

To compare the labeling coverage of Bitbow1 and Bitbow2 flies, we again used the TRH-Gal4 fly to constrain labeling in ^~^100 neurons across the whole larva brain (**Fig. 4b**, **4c**). As expected, we found that Bitbow2.1 had the best labeling coverage, as high as 93.8% in the flies that carried two copies of the transgene (**Fig. 4d**). We also found that colorful Bitbow2 labeling could be achieved by direct crosses to various subtype specific enhancer-Gal4 driver lines (**Fig. 5e** and **Fig. S10**). Finally, when crossing a Bitbow2.1 fly to an elav-Gal4 fly, its offspring labeled neighboring neurons labeled in many distinct Bitbow colors, which indicates that Flp recombination is specific in postmitotic neurons (**Fig. 5f**). Again, the diverse Bitbow colors indicated the srFlp activity is strong yet transient to ensure stable recombination outcomes.

**Figure 5.**
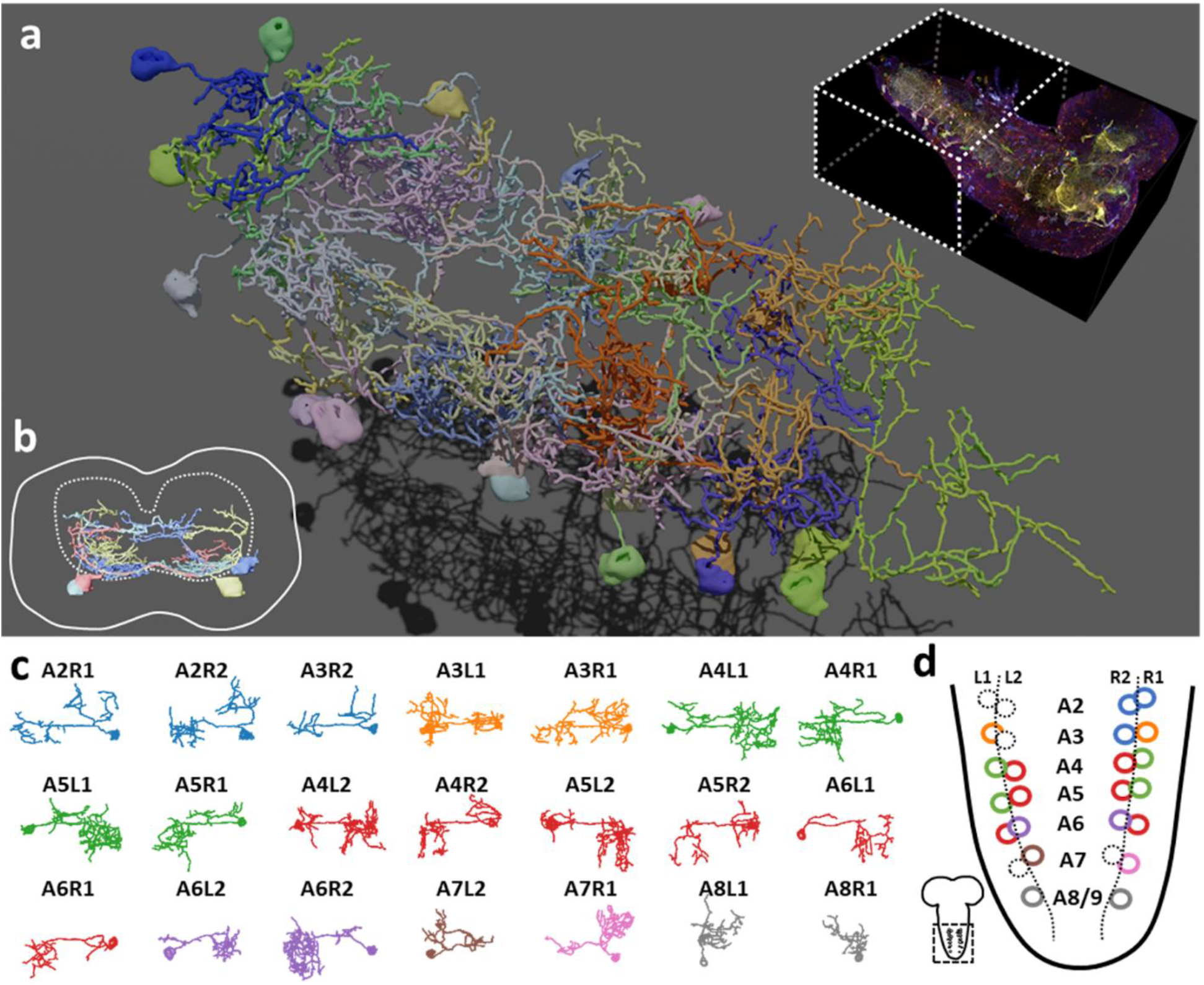
Bitbow2 tracing enables serotonergic neuron morphology and network analysis in larval VNC. **(a)** 3D rendering of traced VNC serotonergic neurons labeled by Bitbow2 in a single brain. Top right, dashed-line box indicates the VNC volume that has been traced. **(b)** Cross-sectional view of the traced serotonergic neurons located in the A5 segment (solid outline), which illustrates that the somas (solid oval shapes) are located at the very ventral part of the VNS while the neurites occupy mostly in the sensory zone (ventral part) of the neuropil (dashed outline). **(c)** Z-projections (dorsal view) of 21 traced serotonergic neurons categorized into 8 morphological subtypes that are indicated by distinct pseudo-colors. **(d)** Schematic of the abdominal VNC and serotonergic neurons. Circles indicate the soma locations of the traced serotonergic neurons, and their colors correspond to the morphological subtype pseudo-colors in (**c**). Dashed circles indicate the soma locations of the unlabeled serotonergic neurons.

### Bitbow2 enables neural anatomy and network analysis in the *Drosophila* central nervous system

As Bitbow2 provides rich color and broad coverage labeling, we expect it can be used to simultaneously resolve many neuron morphologies in the same brain. This not only increases the experimental throughput, but also eliminates the sampling errors and animal-to-animal variations in experiments that rely on aligning sparsely reconstructed neurons from multiple brains to a common reference^53^. To gain higher imaging resolution to resolve the intermingled neurites in the dense neuropil, we applied a modified protein-retention Expansion Microscopy (proExM) protocol to the Bitbow2 *Drosophila* brain (**Fig. S11**)^54–56^. With ^~^4x expansion, we could use nTracer^31^ to reconstruct all 21 Bitbow-labeled (out of 26 estimated total ^46,56^) VNC serotonergic neurons from the A2 to A8/9 segments of a single 3rd instar larva brain (**Fig. 5a** and **Movie S1, S2)**. We sampled the Bitbow colors along the somas and processes of these neurons and found these 21 neurons were labeled by 16 well-separated colors in a UMAP projection (**Fig. S12a**). Although there were 3 unique Bitbow colors, each of which labeled 2, 2, or 4 neurons, their subtle color differences (**Fig. S12b**) and well-separated physical locations (**Fig. S12c**) allowed us to reconstruct their morphology with little ambiguity. We found that all VNC serotonergic neurons project quite locally, mostly within the same segment (**Fig. 5a**). Their somas are located at a very ventral part of the VNC and their projections are mostly restricted to the sensory zone (ventral half) of the VNC (**Fig. 5b**)^57,58^.

As the majority of serotonergic neurons in the A2 to A8/9 segments of this VNC were labeled and reconstructed, we paid extra attention to discover potential anatomical roles that respect the repeated hemi-segment patterns of the VNC (**Fig. S13**). We noticed that all VNC serotonergic neurons within the same hemi-segment send out co-fasciculated neurites that form a single commissure projecting to the contralateral side (**S11e**, arrowheads). While serotonergic neurons in the same hemi-segment have quite distinct morphologies and projection patterns, they have similar counterparts in the contralateral hemi-segment, therefore, forming a bi-lateral symmetric network (**Fig. 5c**, **5d**). These morphologically similar neurons can be classified as at least 8 distinct subtypes based on the quantification of morphological features, including projection density in the contralateral and the ipsilateral side, major branching patterns and anterior *vs* posterior projection distribution (**Fig. 5d**, detailed in **Methods**).

## Discussion

We reported Bitbow, a set of novel transgenic tools capable of generating a large number of unique imaging barcodes in a single animal (**Table 1**). Bitbow utilizes a novel design, in which independent Flp/FRT recombination events lead to binary choices of expressing orthogonal spectral labels. This mechanism exponentially expands the color-coding capacity to 2^N^-1 when using N “bits” of spectrally distinguishable tags. Targeting the same 5-FPs to 3 imaging differentiable subcellular compartments, we created mngBitbow1.0 transgenic flies, which can generate up to 32,767 unique Bitbow codes in a single brain. This is a significant advantage for imaging-based lineage tracing studies because it greatly increases the possibility of labeling neurons with unique lineage codes. Interestingly, we found that heat-shock induced recombination events are constrained in neural stem cells of the larval Bitbow flies. Such serendipity permits directly using Bitbow codes to determine lineage relationships between neural progenies. Providing statistical quantification and modeling, we established that it is feasible to map the lineage relationships between any subtype-specific neurons, driven by any enhancer-Gal4, using fewer than ^~^10 brains.

**Table 1.** Transgenic *Drosophila* Bitbow summary.

| Bitbow version | Subcellular compartment | Insertion site (Chr.) | self-regulating Flp | Labeling density |
| --- | --- | --- | --- | --- |
| **mBitbow 1.0** | Cell membrane | attP40(2L) or attP2(3L) | No | Variable (with hsFlp) |
| **nBitbow 1.0** | Nucleus | attP40(2L) or VK27(3R) | No | Variable (with hsFlp) |
| **gBitbow 1.0** | Golgi | attP40(2L) or attP2(3L) | No | Variable (with hsFlp) |
| **mBitbow 2.0** | Cell membrane | attP2(3L) or VK27(3R) | Yes | Medium (40%-60%) |
| **mBitbow 2.1** | Cell membrane | attP2(3L) or VK27(3R) | Yes | High (70%-95%) |

In practice, we found that certain FP expressed with much higher frequencies than other ones. We suspect that this is due to more frequently spinning of their flanking FRT sites under suboptimal recombination conditions, such as heat-shock induced transient Flp activity. Bitbow codes containing these FPs would have higher collision rates when used in lineage mapping studies. We mitigate such disadvantage by excluding cells labeled by the high-frequency Bitbow codes from analysis. In the future, this problem can be avoided by screening more incompatible FRT sites and using only those with similar recombination efficiencies. Nonetheless, there are other disadvantages associated with heat-shock induced Flp recombination, especially for neuronal morphology labeling and reconstruction. As only the membrane Bitbow is suitable for morphology studies, we found a low percentage of cells were being labeled and their colors were relatively simple that neurons labeled by more than two FPs were relatively rare.

To solve the above-mentioned problems for morphology labeling and reconstruction, we generated Bitbow2 transgenic flies, in which a novel self-regulating Flp module was integrated to effectively recombine mBitbow1 without the need of heat-shock. The elimination of the *hs-* Flp allele yielded two additional advantages: 1) Needing only a simple cross to the broadly used Gal4 libraries, Bitbow2 can be used as a drop-in replacement to any UAS-FP reporters. 2) Abolishing the need for heat-shock, Bitbow2 is compatible with temperature-sensitive assays, such as heat-induced neuronal manipulations with shibire^ts 50^, dTrpA1 ^51^, etc. Finally, we generated different versions of Bitbow2 flies, each of which labeled a different percentage of total neurons to suit the need of tuning the labeling density for different Gal4 driver lines.

Combining sample expansion (ExM) and saturated neuron tracing (nTracer), Bitbow2 flies are suitable for high-throughput morphology studies from light microscopy images. We found that Bitbow labeling is statistically consistent throughout the neuron soma and neurites. This builds the confidence of using fluorescence intensity difference in each spectral channel to differentiate neighboring neurons when using nTracer to reconstruct their morphology. We estimate that thousands (^~^5^5^) of Bitbow “colors” can be easily distinguished in a well-taken 16-bit image dataset. Densely packed neurons, such as the VNC serotonergic neurons are now readily traceable to not only classify the morphological heterogeneity, but also reveal the neural network patterns among a genetically defined population. We envision that, in the future, combining with other high-throughput modalities, such as light-sheet microscopy and automated neuronal tracing, will make larger scale, multi-brain morphological studies feasible in most laboratories.

## Materials and Methods

### Key Resources

| REAGENT or RESOURCE | SOURCE | IDENTIFIER |
| --- | --- | --- |
| **Antibodies** |  |  |
| Rat Anti-mTFP | Cai Lab |  |
| Chicken Anti-GFP | Cai Lab |  |
| Rabbit Anti-mNeonGreen | Cai Lab |  |
| Mouse Anti-mKusabira-Orange2 | Cai Lab |  |
| Guinea Pig Anti-mKate2 | Cai Lab |  |
| Alexa Fluor 594 Donkey Anti-Rat | Life Technologies | A21209 |
| Alexa Fluor 488 Donkey Anti-Chicken | Jackson ImmunoResearch | 703-545-155 |
| Atto 490LS Goat Anti-Rabbit | Hypermol | 2309 |
| CF 555 Donkey Anti-Mouse | Sigma | SAB4600060 |
| CF 633 Donkey Anti-Guinea Pig | Sigma | SAB4600129 |
| Alexa Fluor 647 Donkey Anti-Guinea Pig | Jackson ImmunoResearch | 706-605-148 |
| **Bacterial and Virus Strains** |  |  |
| Mach1-T1 Chemically Competent E. coli | Thermo Scientific | C862003 |
| Stbl3 Chemically Competent E. coli | Thermo Scientific | C737303 |
| **Chemicals, Peptides, and Recombinant Proteins** |  |  |
| Acrylic acid N-hydroxysuccinimide ester | Sigma | A8060 |
| Sodium Acrylate | Sigma | 408220 |
| Acrylamide | Sigma | A9099 |
| N,N'-methylenebisacrylamide | Sigma | M7279 |
| Tetramethylethylenediamine (TEMED) | Sigma | T7024 |
| Ammonium Persulfate | Sigma | A3678 |
| 4-hydroxy-2,2,6,6-tetramethylpiperidin-1-oxyl (4HT) | Sigma | 176141 |
| Startingblock (PBS) blocking buffer | Thermo Scientific | 37578 |
| Proteinase K | NEB | P8107 |
| **Experimental Models: Cell Lines** |  |  |
| D. melanogaster: S2 | Bing Ye Lab |  |
| M. musculus: Neuro-2a | ATCC | CCL-131 |
| **Experimental Models: Organisms/Strains** |  |  |
| D. melanogaster: w1118;; | BDSC | RRID: BDSC_5905 |
| D. melanogaster: hsFlp112;; | BDSC |  |
| D. melanogaster:<br>hsFlp112;Sp/CyO;TM2,Ubx/TM6B,Tb | Bing Ye Lab |  |
| D. melanogaster: hsFlp112;;elav-Gal4 | This study |  |
| D. melanogaster: hsFlp112;TRH-Gal4; | This study |  |
| D. melanogaster: yw;;mBitbow1.0 | This study |  |
| D. melanogaster: yw;;nBitbow1.0 | This study |  |
| D. melanogaster: yw;gBitbow1.0; | This study |  |
| D. melanogaster:<br>yw;gBitbow1.0/CyO;mBitbow1.0,nBitbow<br>1.0 | This study |  |
| D. melanogaster: yw;;mBitbow2.0 | This study |  |
| D. melanogaster: yw;;mBitbow2.1 | This study |  |
| D. melanogaster: yw;;2x[mBitbow2.0] | This study |  |
| D. melanogaster: yw;;2x[mBitbow2.1] | This study |  |
| D. melanogaster: w;elav-Gal4/CyO; | BDSC | 8765 |
| D. melanogaster: w;TRH-Gal4; | BDSC | 38388 |
| D. melanogaster: w;;R53C10-Gal4 | BDSC | 38873 |
| D. melanogaster: w;;R67A06-Gal4 | BDSC | 39397 |
| **Recombinant DNA** |  |  |
| pmAmetrine-N1 | Addgene | 54505 |
| pmTFP1-N1 | Addgene | 54521 |
| pmNeonGreen-N1 | Allele Biotechnology |  |
| pmKusabira-Orange2-N1 | 38 |  |
| ptdKatushka2-N | 39 |  |
| pmCitrine-N1 | Addgene | 54594 |
| pmCherry-N1 | 59 |  |
| pJFRC-MUH | Addgene | 26213 |
| pattB-synaptobrevin-GAL4-hsp70 | Addgene | 46107 |
| pJFRC81-10XUAS-IVS-Syn21-GFP-p10 | Addgene | 36432 |
| pMT-Gal4 | DGRC | 1042 |
| pUAST-Flp | DGRC | 1020 |
| pCAG-FlpINT | Dawen Cai Lab |  |
| pDC-MUH | This study |  |
| pDC-MUH-p10pA | This study |  |
| pCMV-2xmAmetrine-N | This study |  |
| pDC-UAS-mBitbow1.0 | This study |  |
| pDC-UAS-nBitbow1.0 | This study |  |
| pDC-UAS-gBitbow1.0 | This study |  |
| pDC-UAS-mBitbow2.0 | This study |  |
| pDC-UAS-mBitbow2.1 | This study |  |
| pAc5-Flp-p10pA | This study |  |
| pCMV-3FRT-mCherry-F14-mTFP | This study |  |
| pCMV-3FRT-mCherry-F15-mCit | This study |  |
| pCMV-3FRT-F14-mTFP-F15-mCit | This study |  |
| pCMV-3FRT-F14-F15-mCit-F13-mCherry | This study |  |
| pCAG-Flpbow3 | 10 |  |
| **Software and Algorithms** |  |  |
| Fiji | 60 | https://fiji.sc/ |
| nTracer (Fiji plugin) | 31 | https://www.cai-lab.org/ntracer-tutorial |
| Spectral Unmixing (Fiji plugin) | Joachim Walter | https://imagej.nih.gov/ij/plugins/spectral-unmixing.html |
| Lasergene | DNASTAR | https://www.dnastar.com/ |
| Graphpad Prism | Graphpad Software | https://www.graphpad.com/ |
| Vaa3D | 61 | http://home.penglab.com/proj/vaa3d/home/ |
| Blender | Blender Foundation | https://www.blender.org/ |
| NumPy | NumPy.org | https://numpy.org/ |
| umap-learn | 62 | https://github.com/lmcinnes/umap |

### *Drosophila* husbandry

Flies were reared at 25°C on standard CT medium with a 12h/12h light/dark cycle. For heat-shock induced Bitbow labeling experiments, hsFlp;;elav-Gal4 or hsFlp;TRH-Gal4; females were crossed to Bitbow1.0 males, and timed-egg-lay was conducted to collect embryos for the desired time window in vials; afterwards, the vials were placed in a 37°C metal-bead bath for 30min to induce the heat-shock, and kept at 18°C to incubate until ready for dissection.

### Molecular cloning and fly transgenics

To test out new incompatible FRT sites, a series of FRT-FP plasmids were constructed (**Fig. S1**) using the mammalian expression backbone pCMV-N1 (Clontech). FRT-F13, FRT-F14 and FRT-F15 sequences were obtained from a previous study ^34^, and introduced to the following plasmids through PCRs. To test the incompatibility of FRT-F14 and FRT-F15 to three known FRTs (FRT-F3, FRT-5T2, FRT-545), pCMV-3FRT-mCherry-F14-mTFP and pCMV-3FRT-mCherry-F15-mCit were built through sequential assembly of the three FRTs from pCAG-Flybow (Cai lab), mCherry-SV40pA from pmCherry-N1 (Addgene), and F14-mTFP-SV40pA or F15-mCitrine-SV40pA from pmTFP-N1 or pmCit-N1 (Addgene), respectively, using PCR, restriction digestion, and ligation. After the incompatibility test was done, mCherry-pA was removed by digestion with two flanking blunt-end sites (PmlI, EcoRV), and re-ligation to produce pCMV-3FRT-F14-mTFP and pCMV-3FRT-F15-mCit, as the control plasmids. Similar cloning approaches were applied in the next steps. To test incompatibility of FRT-F15 to the other four FRT sites, F15-mCit-pA was moved into pCMV-3FRT-F14-mTFP to produce pCMV-3FRT-F14-mTFP-F15-mCit, from which mTFP-SV40pA was removed to produce the control plasmid pCMV-3FRT-F14-F15-mCit. Finally, to test incompatibility of FRT-F13 to the other five FRT sites, F13-mCherry-pA was moved into pCMV-3FRT-F14-F15-mCit to produce pCMV-3FRT-F14-F15-mCit-F13-mCherry, from which mCitrine-SV40pA was removed to produce the control plasmid pCMV-3FRT-F14-F15-F13-mCherry.

For Bitbow1 plasmids, cDNAs encoding the following FPs were used: mAmetrine, mTFP, mNeonGreen, mKusabira-Orange2, and tdKatushka2 ^35–39^. *Drosophila* myristoylation signal peptide of dSrc64B (1-10aa, dMyr), Human histone 2B protein (full length, hH2B) or Mouse Mannosidase II alpha 1 (1-112aa, mManII) was fused in-frame to the N-terminus of individual FPs (dMyr-FP, hH2B-FP, mMannII-FP), to achieve targeted labeling at the cell membrane, nucleus or Golgi apparatus ^63–65^. Individual incompatible FRT sequence pairs (FRT-F13, FRT-F14, FRT-545, FRT-F3, or FRT-5T2) ^10,34,66–68^ were then placed in the opposing direction to flank dMyr-FP / hH2B-FP / mMannII-FP sequence. An upstream activation sequence (UAS) and a p10 polyadenylation sequence (p10pA) ^69^ were placed upstream and downstream of each FRT flanked FP cassettes, respectively, and separately cloned into pDC-MUH, which was based on the pJFRC-MUH backbone vector^70^ with a few digestion site modifications, by standard cloning methods. The Bitbow1.0 plasmids were finally assembled together from the five individual modules through Gibson assembly ^71^.

For Bitbow2 plasmids, the nSyb-promoter-driving self-regulating flippase module was constructed by flanking FlpINT (flippase with an inserted c. elegans intron ^72^, Cai lab) cDNA with a FRT-F14 pair or a FRT-F15 pair which were oriented in the same direction, and then placed downstream of a *Drosophila* n-Synaptobrevin promoter ^73^. The module was then inserted into the mBitbow1.0 plasmid, at a location far away from all 5 FP modules, through Gibson Assembly to generate mBitbow2.0 or mBitbow2.1.

The final Bitbow plasmids were integrated into *Drosophila melanogaster* genome docking sites attP40, attP2 or VK00027 (**Table 1**) through ΦC31-integrase-mediated transgenesis ^74–78^. Embryo injections and transgenic selections were done by BestGene Inc (Chino Hills, CA).

### Dissection and mounting

Adult or 3^rd^ instar *Drosophila* brains were dissected in PBS at room temperature (abbr. RT) within 30min before proceeding to fixation. Dissected brains were fixed in 4% PFA (Sigma #P6148, diluted in PBS) at RT with gentle nutation for 20min, followed by three quick PBST (PBS+1% Triton X-100) washes, then PBS washes for 15min x 3. Brains then either proceeded to direct mounting (for native fluorescence imaging) or immuno-stainings. Vectashield (Vector Laboratories, H-1000) was used as the mounting medium.

### Immunohistochemistry

Fixed brain samples were treated with StartingBlock (Thermo, 37578) for 1 hour at RT with gentle nutation. After blocking, the brains were incubated with primary antibodies diluted in StartingBlock for 2 overnights at 4°C. Three quick PBST washes and PBS washes for 15min x 3 were done, before the brains were incubated with secondary antibodies diluted in StartingBlock for 2 overnights at 4°C. Finally three quick PBST washes and PBS washes for 15min x 3 were done and the brains were ready for imaging. For detailed antibody combinations and dilutions see ***Key-resource-table***.

### Expansion Microscopy (ExM)

ExM brain samples were generated following the ProExM protocol ^54^ with modifications. Antibody-stained Bitbow samples were treated in Acrylic acid N-hydroxysuccinimide ester (AaX, Sigma, A8060) at RT for 1 overnight, followed by PBS washes for 15min x 3. Samples were then incubated in the ExM monomer solution (“Stock-X”, containing Acrylate, Acrylamide, and Bis-acrylamide) at 4°C for 1 overnight. Samples were transferred to fresh ExM monomer solution with gel initiators (APS, TEMED, 4-HT) at 4°C for 15min, and then quickly mounted on a sample chamber made with 200μm adaptors (Sun lab) on a glass slide, sealed with a 22×30 coverslip on top (Fisher, 12-544). The slide was then transferred to a humidity box and incubated at 37°C for about 2 hours until the gel fully polymerized. The gel was trimmed carefully with a razor to allow as little of excessive space around the brains as possible. Trimmed gel pieces were transferred to an EP tube and digested with Proteinase K (NEB, P8107) at 37°C for 1 hour. Three quick PBST washes and PBS washes for 15min x 3 were done before the brains were put into another round of antibody staining, following the same IHC protocol mentioned above. After the second-round staining, the gels were slowly expanded to the final size by changing the submerging solution from PBS to pure diH2O, and ready for imaging.

### Confocal microscopy and linear unmixing

Confocal images were acquired with Zeiss LSM780 with a 20x 1.0 NA water immersion objective (421452-9800-000) or a 40x 1.3 NA oil immersion objective (421762-9900-000). The 32-channel GaAsP array detector was used to allow multi-track detection of five fluorophores with proper channel collection setups (**Fig. S2**).

Spectral Unmixing plug-in (by Joachim Walter) in Fiji was used to perform linear unmixing on Bitbow images. Reference unmixing matrix was measured by imaging cultured mouse N2A cells expressing mAmetrine, mTFP, mNeonGreen, mKO2 or tdKatushka2 separately, with the exact same multi-track setups intended for Bitbow brains. Customized ImageJ scripts were used to automate the unmixing process as well as creating composite image stacks from unmixed channels.

### Image stitching and neuron tracing

When the region of interest was larger than the objective field of view, multiple confocal tiles were taken and stitched offline. 5% overlapping seams were set between adjacent tiles to allow reliable stitching and maximize the area of coverage. Alignmaster 1.0.6 (part of the nTracer tool set) was used to perform stitching between tiles sequentially.

All neuronal tracings were done using nTracer 1.3.5 ^31^. Sampling tolerance for color and intensity were set at 0.3 to allow accurate and efficient tracings. Tracing results were exported in SWC format for downstream 3D-rendering and Bitbow color analysis.

3D visualizations of neuron tracings were performed using custom scripts in the open-source modeling software Blender v2.81 (Blender Foundation). Models containing fluorescence data were produced with a modification of the method described in ^79^.

### Quantification and statistical analysis

#### Bitbow code quantifications

Bitbow-labeled neural clusters in the 3rd instar larval brains were used to quantify the labeling performance of 1-localization, 2-localization, and 3-localization Bitbows. Clusters in the central brain, gnathal segments, and thoracic segments were marked with the Fiji ROI tool, and the on/off status of each color channel in every cluster was manually recorded as 1/0 for each “bit” (examples in **Fig. 1e, 2f**). The frequency of occurrence of each Bitbow module was calculated in each brain, and summarized across multiple brains with the mean and standard deviation of the frequency reported (**Fig. 1h, S7a**). 15-module frequencies of mngBitbow were used to generate empirical probability distribution of all 32,767 mngBitbow codes, which was further used in simulations described in **Fig. 2g, 2j** (details below).

#### Bitbow color differentiation and stability analysis

Pixel intensity values from 5 channels along the tracing of all 21 somas and part of 4 neurites (A5L1, A5L2, A5R1, A5R2) were used to generate analysis on differentiation power as well as the stability of Bibtow labeling. Raw intensities were processed through a 3×3×3 median kernel and a 10-pixel rolling window average to reduce noise, then the pixel intensity in each channel was normalized to the sum of five channels of that pixel, in order to bring brighter and dimmer pixels to the same scale for accurate color analysis.

To visualize the differentiation power using all five Bitbow channels, UMAP was used on the 21 somas to reduce the five-dimension pixel values onto a 2D embedding. To visualize differences between 4 relatively closely-located neurons on the 21-soma embedding, an independent UMAP was generated with only those 4 neurons using the same parameters. Data were processed with Python 3.7.4 and umap-learn 0.3.10.

To visualize the stability of Bitbow labeling, soma pixel intensities and neurite pixel intensities from the same neurons were summarized in “split-violin” plots, where in each plot the left half represents soma pixels and the right half neurite pixels.

#### Simulations of Bitbow labeling requirements

Computer simulations were used to estimate the number of animals required to achieve saturated coverage for a range of hypothetical N-neuron systems (**Fig. 2j**). For each condition, 500 trials of Algorithm 1 were averaged using custom code implemented in Python v3.7.4 and NumPy v1.17.2. Here, *activation_rate* was assigned to estimates of 0.5% (MARCM) and 48.08% (Bitbow) to model differences in labeling densities. The whitelist array *valid_barcodes* was assigned as the k lowest-probability barcodes (32,767, 32,700 or 32,000) for the Bitbow trials or all possible barcodes for MARCM.

**Algorithm 1:**
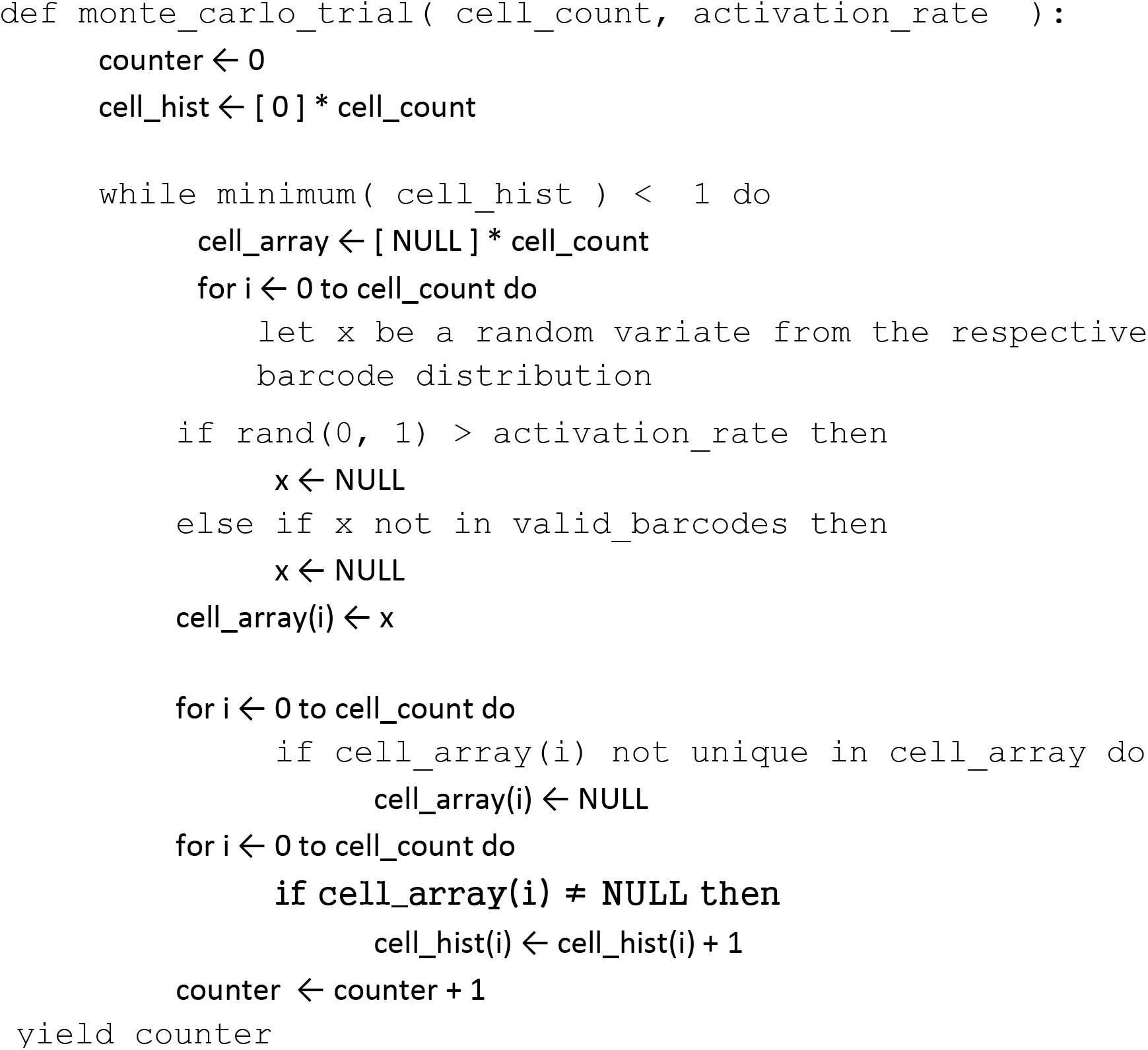

#### Simulations of Bitbow barcode collision rate

Simulations of the empirical collision rate (**Fig. 2g**) were performed in a similar manner. Briefly, random lists of barcodes were drawn from the empirical distributions, followed by the counting of repeated barcodes to produce an overlap rate. This process was repeated 100,000 times for systems under 100 lineages, 1,000 times for systems between 100 and 1000 lineages, and 10 times between 1,000 and 1,000,000 lineages, due to computational complexity.

## Data Availability

The data that support the findings of this study are available from the corresponding author upon reasonable request.

## Code Availability

Custom code for analysis of image processing and data analysis is available from the corresponding author upon reasonable request.

## Acknowledgments

We thank all members of the Cai lab who contribute to the discussion and revision of the manuscript. We thank Bing Ye for providing several transgenic *Drosophila* lines. We thank Grace Hyunh and Ed Boyden for their advice on expansion microscopy. YL acknowledges support by the Patten Fellowship. MG acknowledges support by the University of Michigan UROP summer fellowship. DC acknowledges support by NIH 1R01AI130303, 1UF1NS107659, 1R01MH110932, and 1RF1MH120005, and by National Science Foundation NeuroNex-1707316.

## Author Contributions

DC conceived and supervised the project. YL and DC designed the experiments. YL, LAW and DC wrote the manuscript with input from all authors. YL designed cloning strategies to construct Bitbow plasmids. YL, YZ, MG, TYC and DHR generated the Bitbow transgenic flies. YL, YZ, and EME processed brain samples and performed microscopy. YL and MC quantified the Bitbow1 lineage codes. YL and EME quantified Bitbow2 labeling coverage. YL and HPJC traced the VNC serotonergic neurons. LAW, YL, NSM and DC established the statistical models. YL wrote the codes for Bitbow color analysis. LAW wrote the codes for simulations and scripts for 3D renderings of traced neurons.

