## Supplemental Figure & Movie Legends for "Bitbow: a digital format of Brainbow enables highly efficient neuronal lineage tracing and morphology reconstruction in single brains"

#### Supplemental Figures

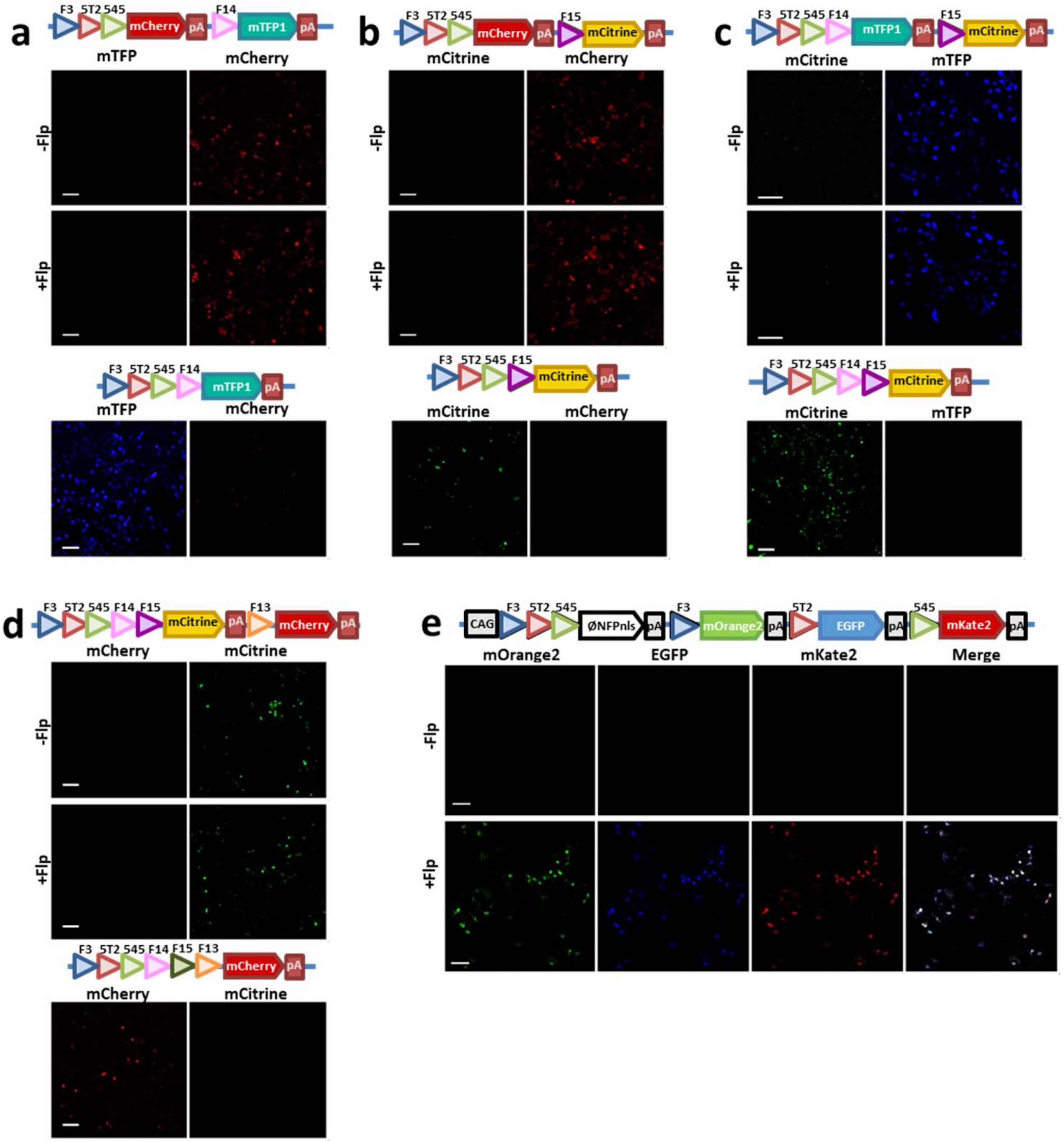
a
F3 5T2 545
mCherry
pA
F14
mTFP1
pA
mTFP
mCherry
-Flp
+Flp
F3 5T2 545 F14
mTFP1
pA
mTFP
mCherry
b
F3 5T2 545
mCherry
pA
F15
mCitrine
pA
mCitrine
mCherry
-Flp
+Flp
F3 5T2 545 F15
mCitrine
pA
mCitrine
mCherry
c
F3 5T2 545 F14
mTFP1
pA
F15
mCitrine
pA
mCitrine
mTFP
-Flp
+Flp
F3 5T2 545 F14 F15
mCitrine
pA
mCitrine
mTFP
d
F3 5T2 545 F14 F15
mCitrine
pA
F13
mCherry
pA
mCherry
mCitrine
-Flp
+Flp
F3 5T2 545 F14 F15 F13
mCherry
pA
mCherry
mCitrine
e
CAG
F3 5T2 545
ØNFPnls
pA
F3
mOrange2
pA
5T2
EGFP
pA
545
mKate2
pA
mOrange2
EGFP
mKate2
Merge
-Flp
+Flp

**Figure S1. Validation of additional orthogonal FRT sites used in the Bitbow constructs.** Series of constructs were generated and co-transfected with pCAG-Flp into mouse N2A cells to determine the orthogonality of FRT variants in addition to the previously reported FRT-F3 (blue triangles), FRT-5T2 (green triangles) and FRT-545 (red triangles) sites (3FRT). **(a, b)** show the results of validation of FRT-F14 (pink triangles) and FRT-F15 (magenta triangles), respectively. In either experiment, mCherry expression was driven by the CMV promoter and the mRNA was terminated by a polyadenylation (pA) sequence positioned right after the mCherry. If the FRT-F14 or FRT-F15 site can be recombined with any of the 3FRT sites, the removal of mCherry-pA sequence would result in the expression of mTFP or mCitrine, respectively. We found that regardless of whether Flp is co-expressed or not, only mCherry expression was observed, which indicated the FRT-F14 or FRT-F15 site is incompatible to the 3FRT sites (top panel of **a** or **b**, respectively). Next, we used restriction enzymes to remove the mCherry-pA sequence and found that mTFP or mCitrine can indeed be properly expressed if this was caused by Flp/FRT recombination (bottom panel of **a** or **b**, respectively). **(c)** To validate whether FRT-F14 and FRT-F15 are incompatible to each other, we took the pCMV-3FRT-F14-mTFP-pA construct from (**a**) and concatenated F15-mCitrine-pA to its 3'-end. We found that regardless whether Flp is co-expressed or not, only mTFP expression was observed, which indicated that FRT-F14 is incompatible to FRT-F15. (**c**, top panel). Similarly, we generated a 5FRT-mCitrine-pA plasmid and confirmed the proper expression of mCitrine if mTFP-pA sequence was removed (**c**, bottom panel). **(d)** To validate whether FRT-F13 is incompatible to the other 5 FRT sites, we took the 5FRT-mCitrine-pA and concatenated F13-mCherry-pA to its 3'-end. We found that regardless whether Flp is co-expressed or not, only mCitrine expression was observed, which indicated that FRT-F13 is a 6th incompatible FRT site. (**d**, top panel). And again, we generated a 6FRT-mCherry-pA plasmid and confirmed the proper expression of mCherry if mCitrine-pA sequence was removed (**d**, bottom panel). **(e)** The activity of Flp in experiments (**a-d**) was demonstrated by including a control, in which the pCAG-Flpbow3 plasmid (Cai *et al.* 2013) was co-transfected with or without a pCAG-Flp plasmid.

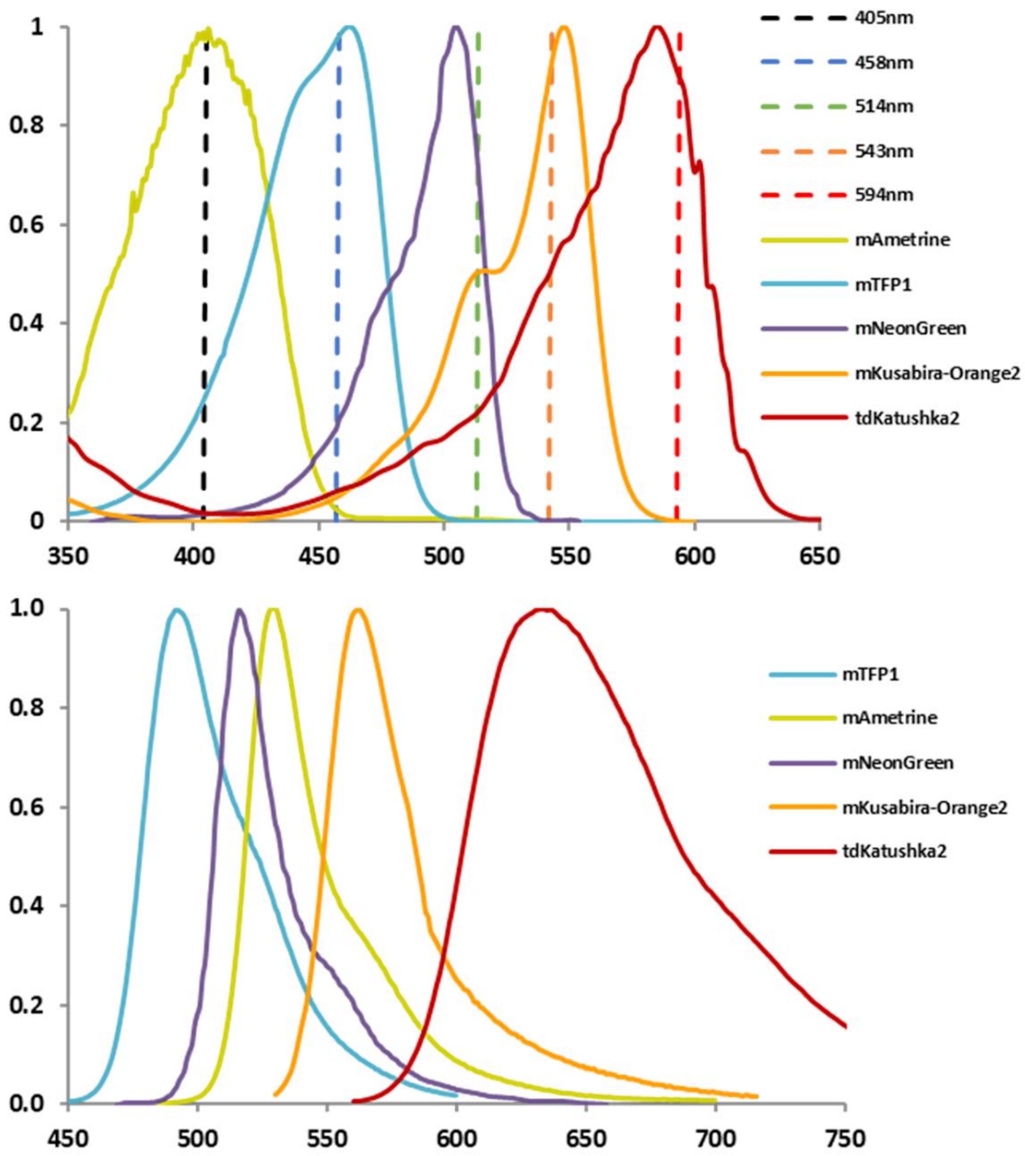


| Track | MBS | Laser | Channels (Fluorophores) |
|---|---|---|---|
| I | -405/458/514/594 | 405nm | 456-501(mTFP*), 501-545(mAme) |
| II | -405/458/514/594 | 458nm | 456-501(mTFP), 501-545(mNG*) |
| III | -405/458/514/594 | 514nm, 594nm | 501-545(mNG), 545-572(mKuO2), 625-696(tdKatu2) |

**Figure S2. Spectral property and confocal imaging setting for the five Bitbow fluorescent proteins (FPs).** Top panel, normalized absorbance (excitation) efficiency spectra of the five FPs. Dashed vertical lines, wavelength of excitation lasers used to image the five FPs. Middle panel, normalized emission efficiency spectra of the five FPs. Bottom panel, confocal microscope setup to sequentially image these FPs in line-switching mode. MBS, main beamsplitter. Asterisks, extra recording channels used for spectral linear unmixing (detailed in **Methods**).

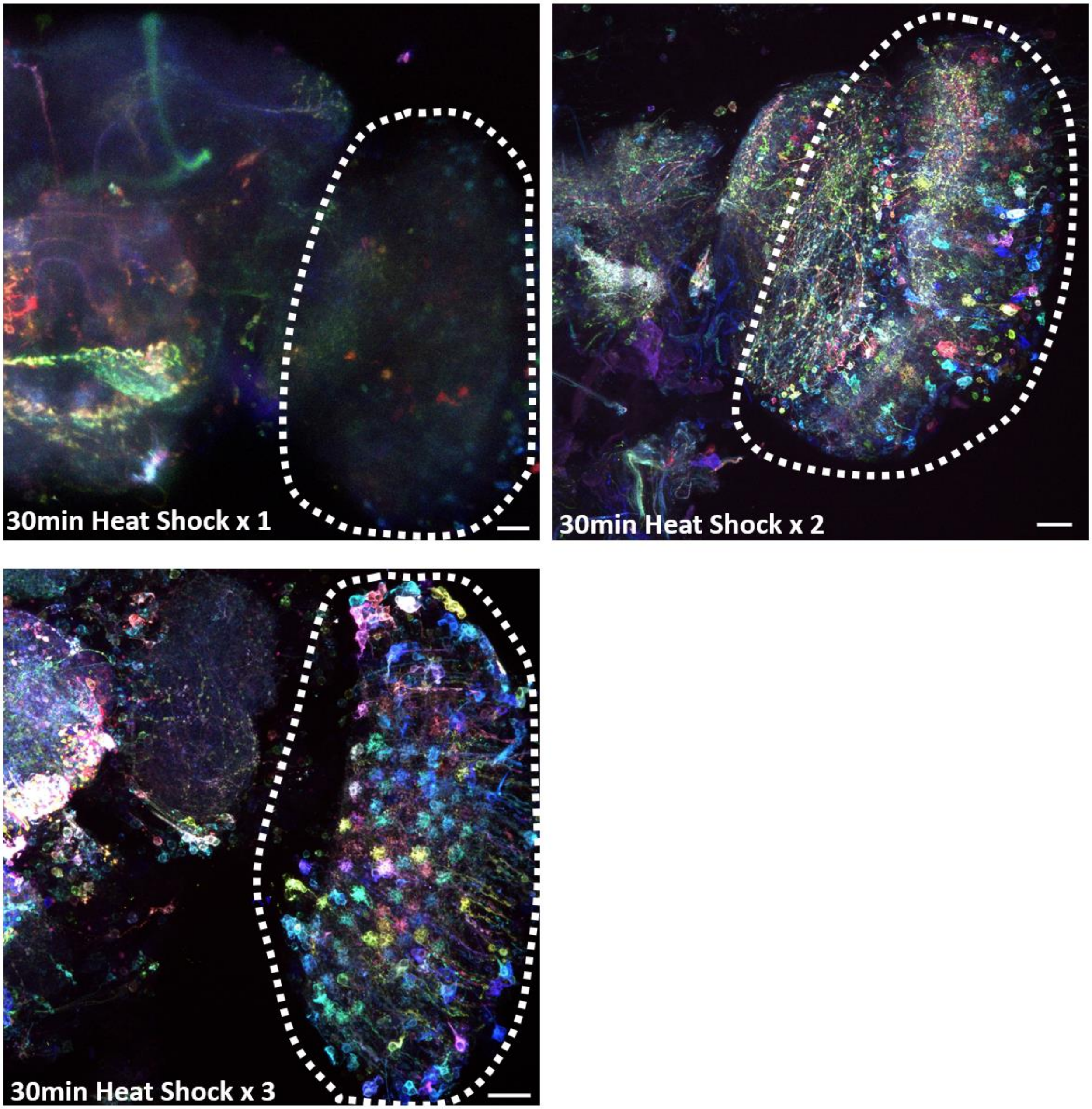


**Figure S3. Multiple heat-shock induced flippase activity labels more neurons in mBitbow1.0 flies.** Adult fly brains from a hsFlp;;elav-Gal4 to mBitbow1.0 cross were experimented with one, two or three 30-minute heat-shocks, similar to the protocol shown in **Fig. 1c**. Dashed regions, optic lobes. Scale bars are 20μm.

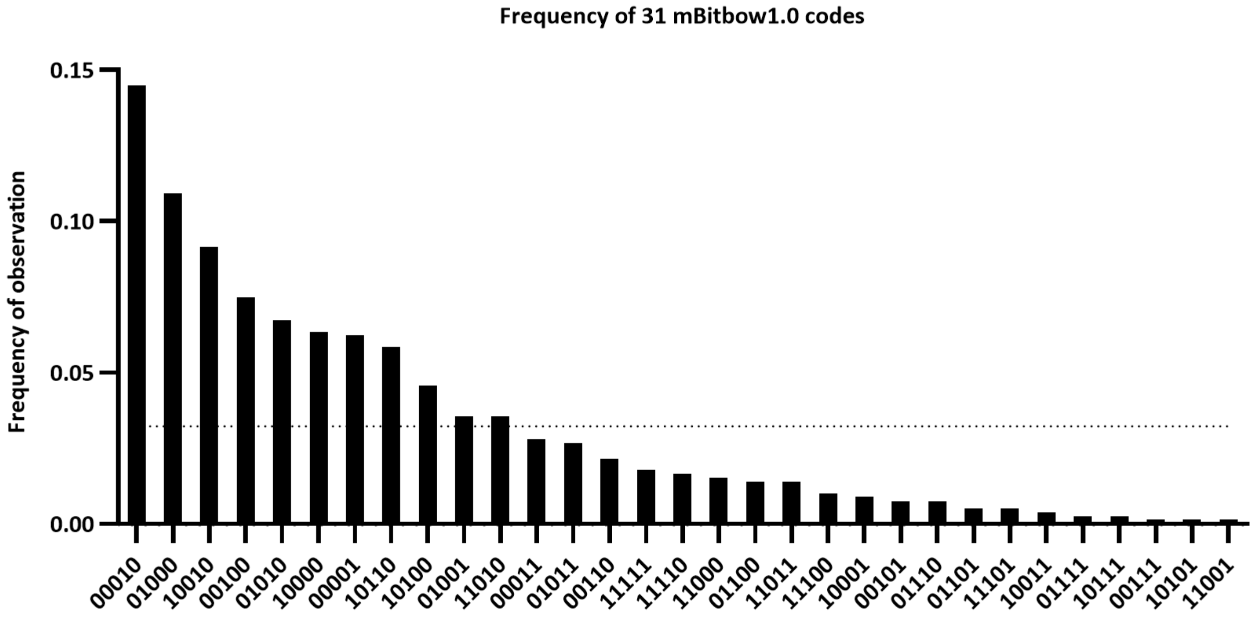


**Figure S4. Observed frequency of the 31 mBitbow1.0 codes upon heat-shock induced Flp recombination.** Bar graph, observed frequency of each Bitbow code. Dotted line, expected frequency if all codes appear in equal chance. N = 787.

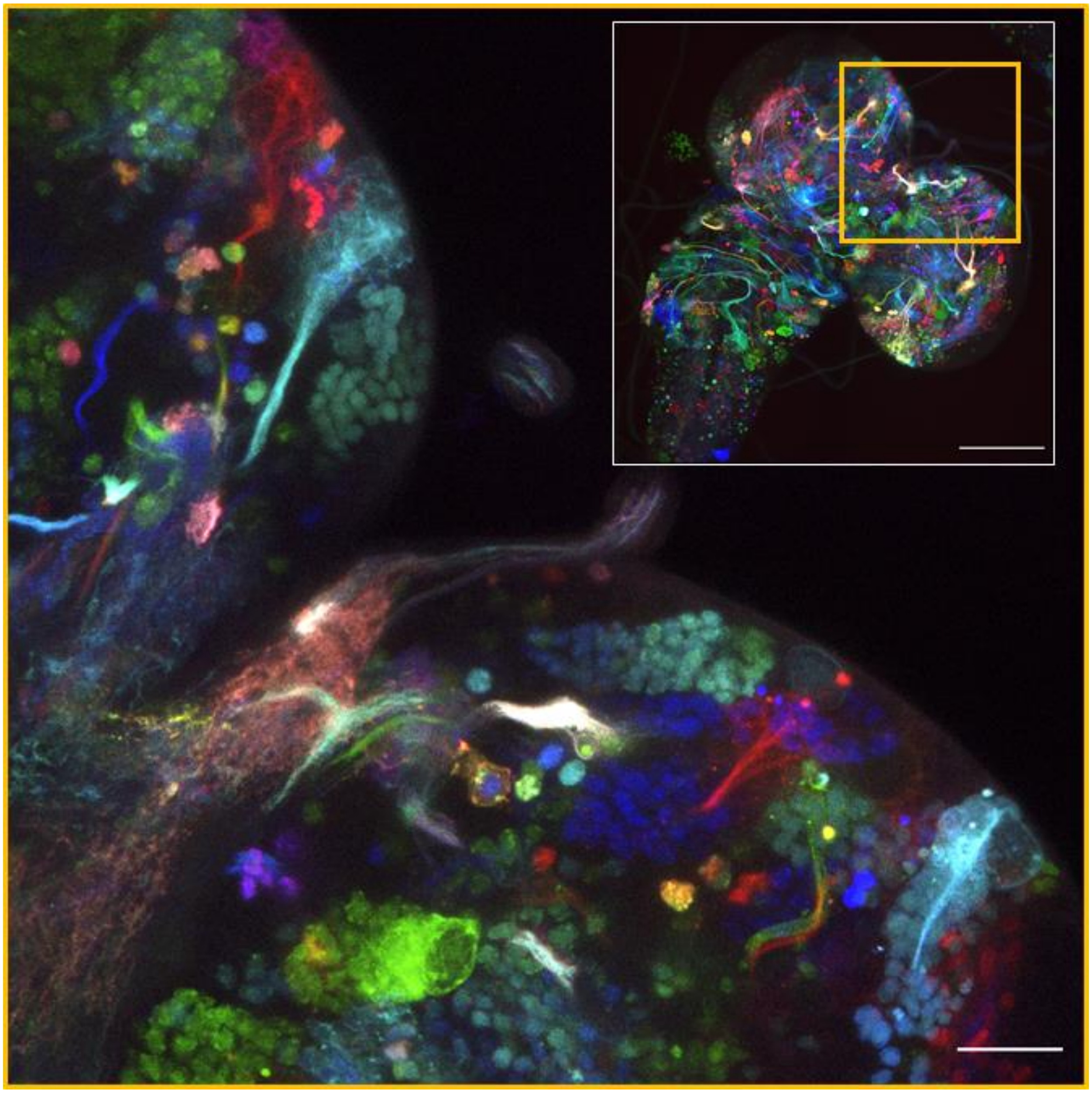

**Figure S5. Third-instar larva neuronal clusters labeled by mnBitbow1.0.** A hsFlp;;elav-Gal4 driver fly was crossed to the membrane and nucleus dual-targeted mnBitbow1.0 fly, whose offsprings were heat-shock at the 3rd instar larva stage, similar to the protocol shown in **Fig. 1f**. The white box shows the maximum intensity projection of the whole brain overview and the yellow boxed region is magnified to show a partial z-plane maximum intensity projection. Scale bars: overview 100µm, magnified 20µm.

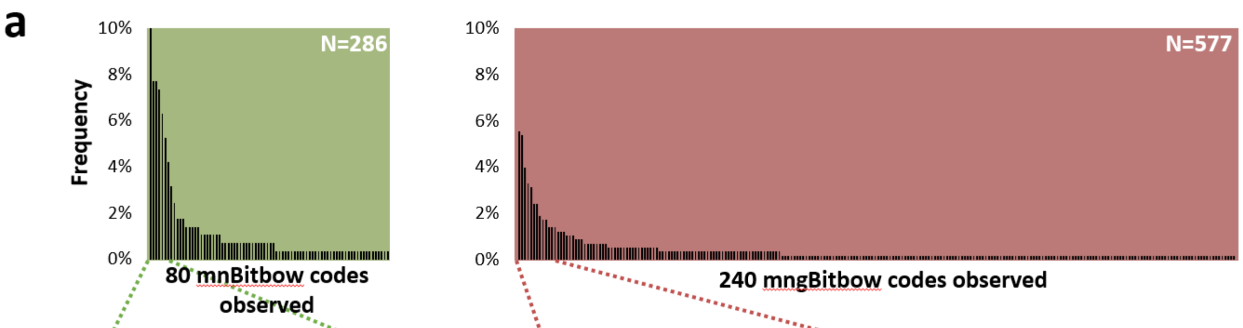


**b**

| Rank | mnBitbow code | Frequency |
|---|---|---|
| 1 | 0000000100 | 10.49% |
| 2 | 0000000110 | 7.69% |
| 3 | 0010000000 | 7.69% |
| 4 | 0000001000 | 7.34% |
| 5 | 0010000100 | 6.29% |
| 6 | 0000001100 | 5.24% |
| 7 | 0000010100 | 4.20% |
| 8 | 0011000000 | 3.15% |
| 9 | 0000100000 | 2.45% |
| 10 | 0000010110 | 1.75% |

| Rank | mngBitbow code | Frequency |
|---|---|---|
| 1 | 000000000000100 | 5.55% |
| 2 | 000000000000001 | 5.37% |
| 3 | 000000010000000 | 3.99% |
| 4 | 001000000000000 | 3.29% |
| 5 | 010000000000000 | 3.12% |
| 6 | 000000000000110 | 2.43% |
| 7 | 001000010000000 | 2.43% |
| 8 | 000000010000100 | 1.91% |
| 9 | 000000100000000 | 1.73% |
| 10 | 001000000000100 | 1.73% |

**Figure S6. Statistics of observed mnBitbow and mngBitbow codes. (a)** Histograms of the mnBitbow or mngBitbow code frequencies observed from brains sampled in **Fig. 2**. **(b)** Top 10 most frequently observed mnBitbow or mngBitbow codes. Bits corresponding to mNeonGreen expressions were highlighted in green.

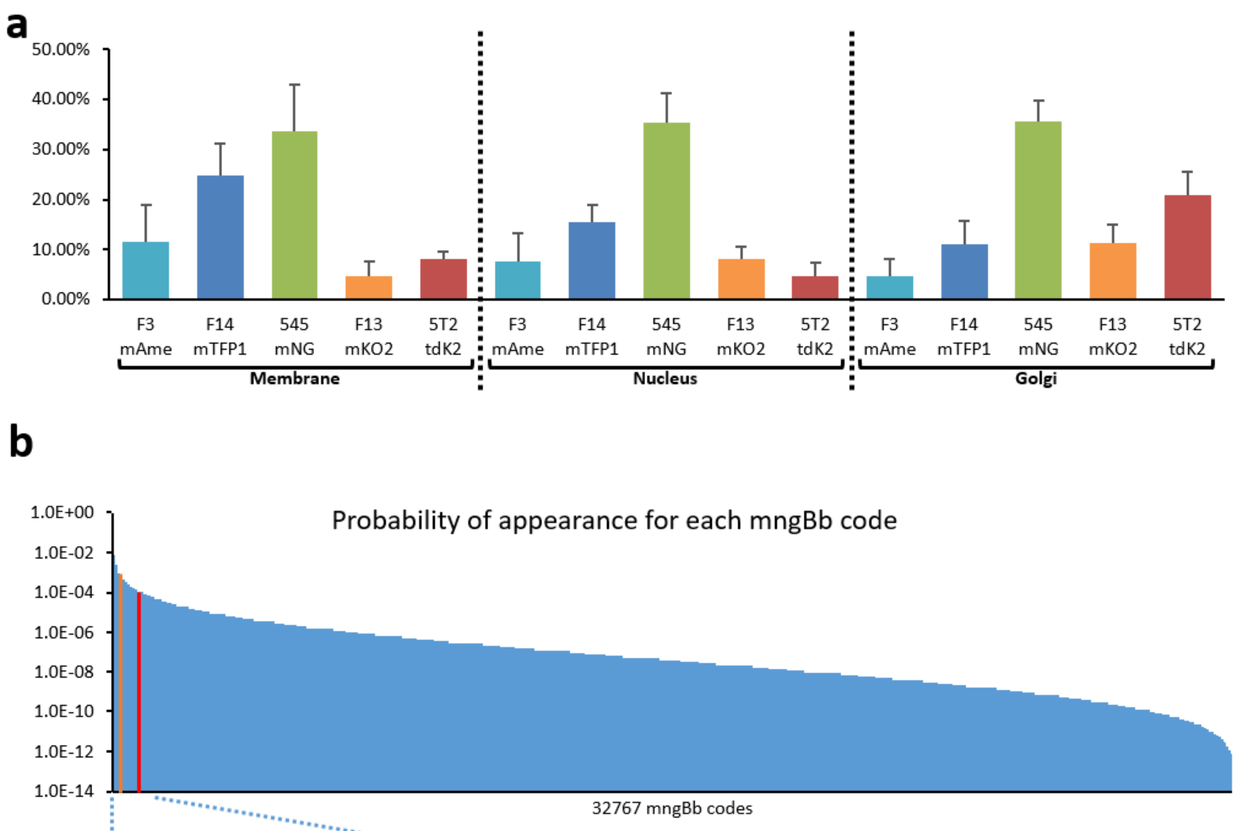


c

| | | | | | | | | | | | | | | | | |
|---|---|---|---|---|---|---|---|---|---|---|---|---|---|---|---|---|
| | 000000000000100 | 000000010000000 | 001000000000000 | 010000000000000 | 000000010000100 | 001000000000100 | 001000010000000 | | 100000010000100 | 000000010000110 | 000000010001100 | | 101010010000100 | 101000011000100 | 001000010100010 | |
| **p-value** | 0.0362 | 0.0360 | 0.0334 | 0.0217 | 0.0198 | 0.0184 | 0.0183 | ... | 2.57E-03 | **2.50E-03** | 2.46E-03 | ... | 1.15E-04 | **1.15E-04** | 1.15E-04 | ... |
| **rank** | 32767 | 32766 | 32765 | 32764 | 32763 | 32762 | 32761 | ... | 32701 | **32700** | 32699 | ... | 32001 | **32000** | 31999 | ... |

**Figure S7. Estimation of the experimental frequencies of mngBitbow1.0 codes. (a)** Observed frequencies of the 15 recombination events in mngBitbow1.0. **(b)** Based on the observed frequencies quantified in (**a**), we synthesized the probability of occurrence of all 32767 mngBitbow codes. **(c)** High-probability mngBitbow codes and their corresponding p-values are sorted from high probability to low probability. Bits corresponding to mNeonGreen expressions were highlighted in green. Code cutoffs at rank 32700 and 32000 used in **Fig. 2g** and **2j** were highlighted in yellow and red, respectively.

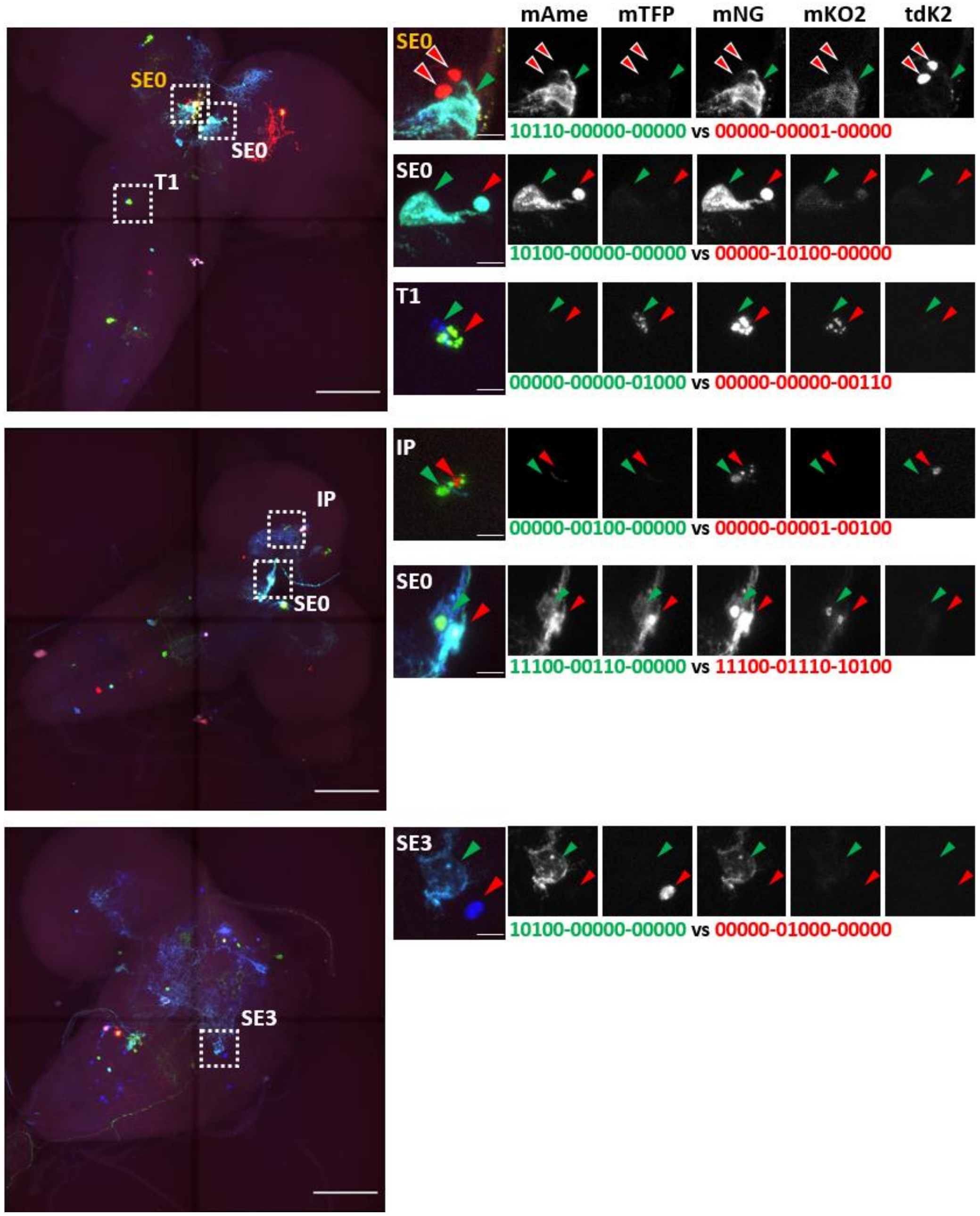


**Figure S8. Examples of determining the lineage relationships between larval serotonergic neuron clusters.** Related to **Fig. 3**, all neuron clusters identified from each brain are magnified from the boxed regions of the overview images. mngBitbow codes in green and red correspond to the neurons marked by the green and red arrowheads, respectively. Among all the observed serotonergic neuron clusters, only two SE0 neurons in one brain (the very top panel) are marked by the same mngBitbow code 00000-00001-00000, which has a relatively high probability of occurrence at 0.00327 and ranked as 32721/32767 among all possible mngBitbow codes (included but not displayed in **Fig. S7b**). Scale bars: overview 100µm, magnified panels, 10µm.

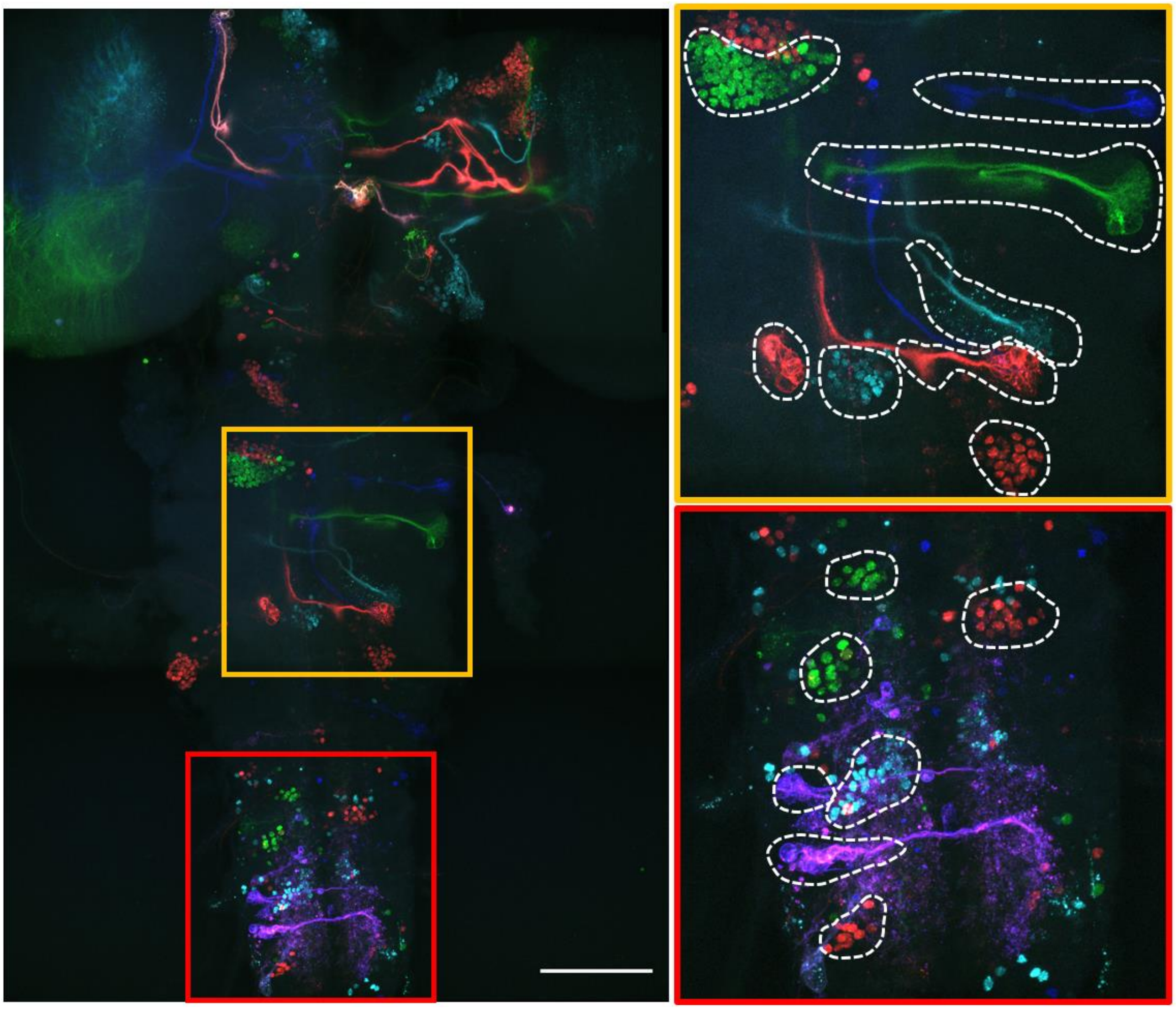

**Figure S9. Early embryonic recombination resulted in many neuronal clusters labeled in the same Bitbow codes.** Left panel, maximum intensity projection of a 3rd instar brain labeled by an early-embryonic (0-4h AEL) heat-shock of the offspring of the mngBitbow1.0 x hsFlp;;elav-Gal4 cross. Right panels, magnified yellow and red boxed regions in the left panel. Dashed outlines indicate individual neuron clusters labeled by the same Bitbow codes. Scale bar, 100µm.

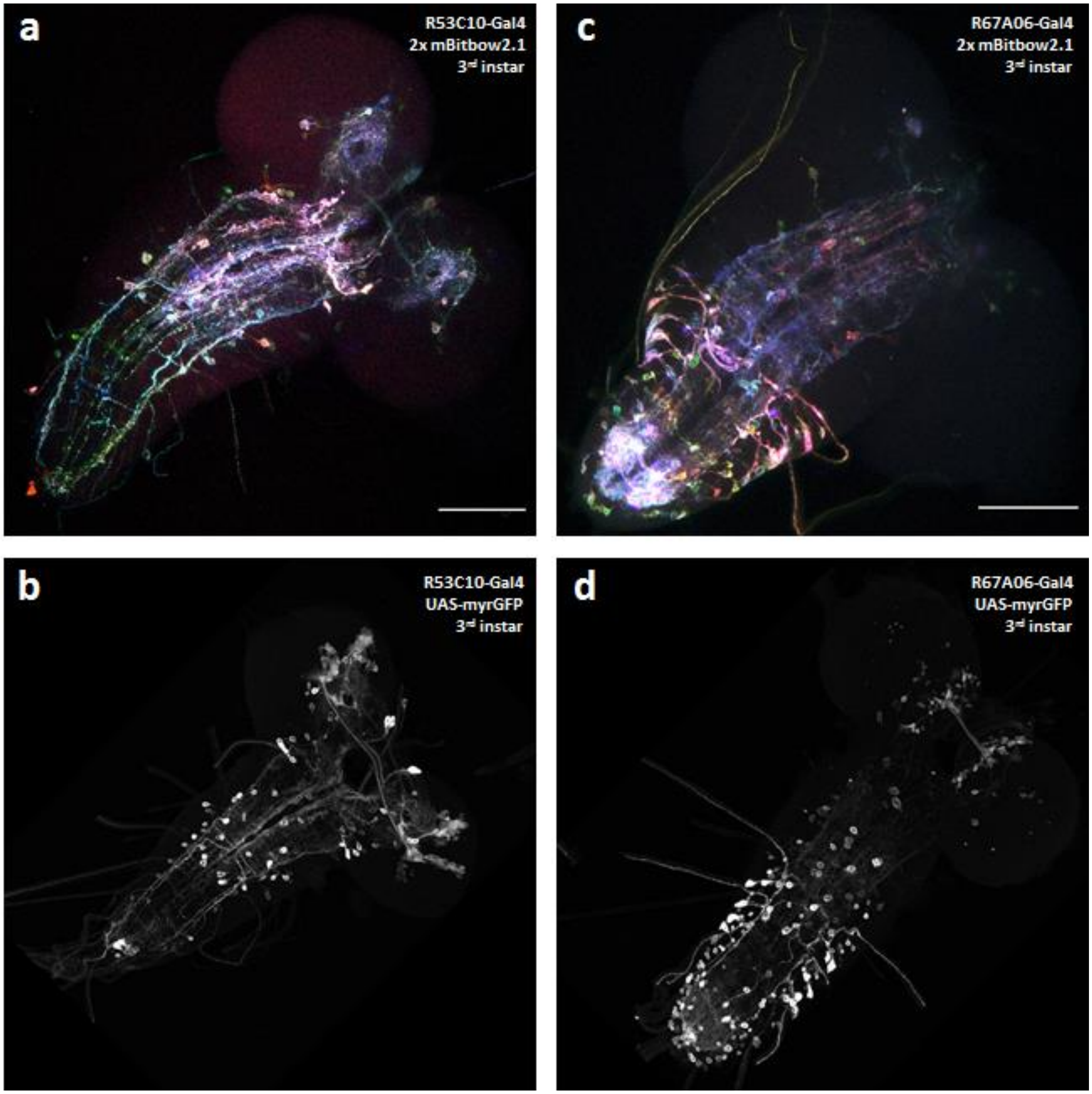


**Figure S10. Bitbow2.1 labels different subsets of neurons when directly crossed to specific enhancer-Gal4 drivers. (a, b)** 3rd instar neurons labeled in an offspring of the 2x mBitbow2.1 fly or UAS-myrGFP (from Janelia FlyLight collection) crossed to the R53C10-Gal4 fly, respectively. (c, d) 3rd instar neurons labeled in an offspring of the 2x mBitbow2.1 fly or UAS-myrGFP (from Janelia FlyLight collection) crossed to the R67A06-Gal4 fly, respectively. Scale bars: 100µm.

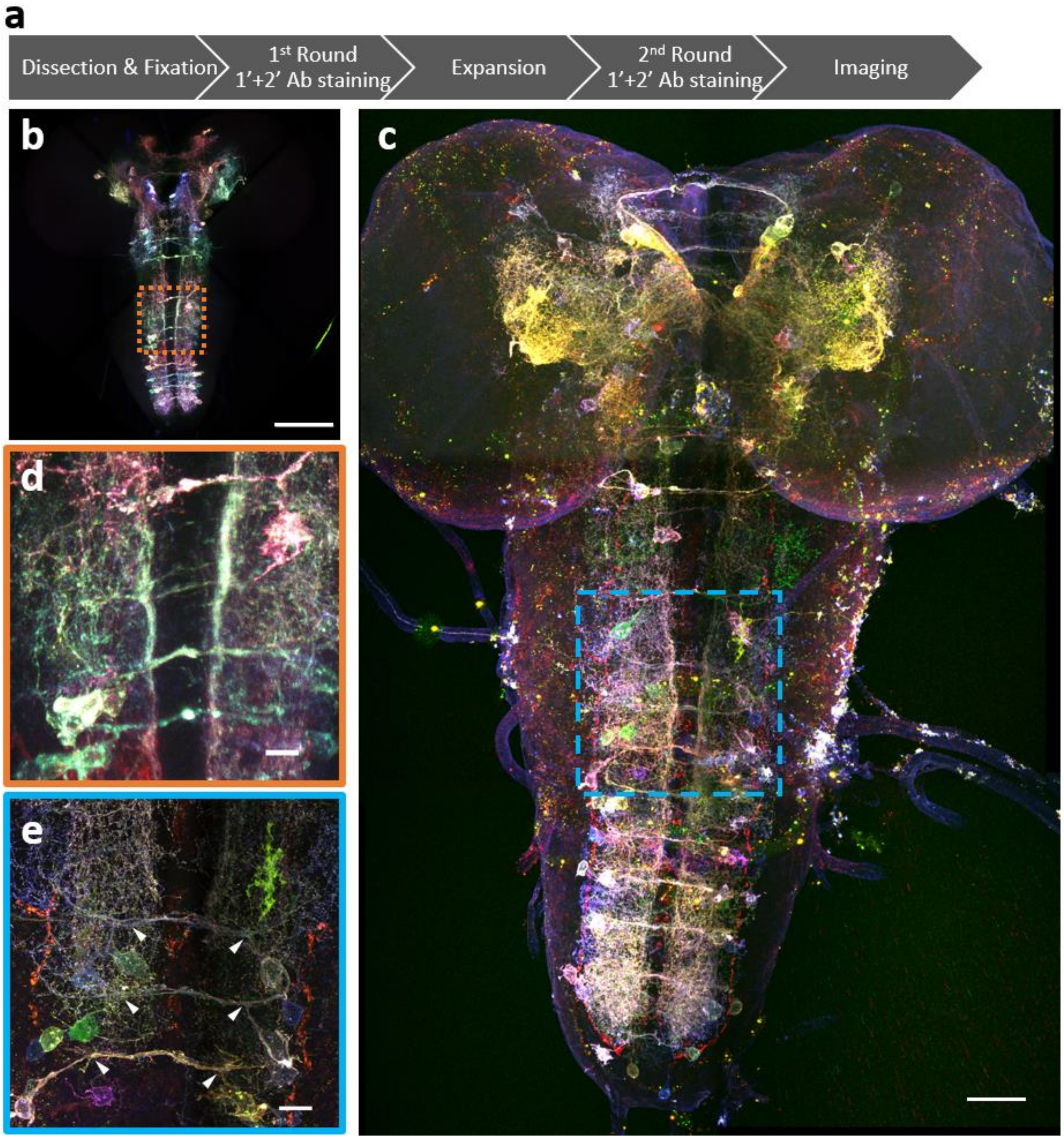


**Figure S11. Super-resolution Bitbow imaging enabled by a modified protein retention-Expansion Microscopy (pro-ExM) protocol. (a)** Experimental flow of the modified pro-ExM protocol. **(b)** Serotonergic neurons labeled by TRH-Gal4 driven 2x Bitbow2.1 without sample expansion and imaged by native fluorescence. **(c)** Serotonergic neurons labeled by TRH-Gal4 driven 2x Bitbow2.1 after ~4x sample expansion and imaged by immuno-fluorescence. **(d, e)** Magnified boxed regions in (**b**, **c**), respectively. Arrowheads indicate VNC serotonergic neurons within the same hemi-segment send out co-fasciculated neurites that form a single commissure projecting to the contralateral side. Scale bars, (**b**, **c**) 100µm, (**d**) 10µm, (**e**) 35µm.

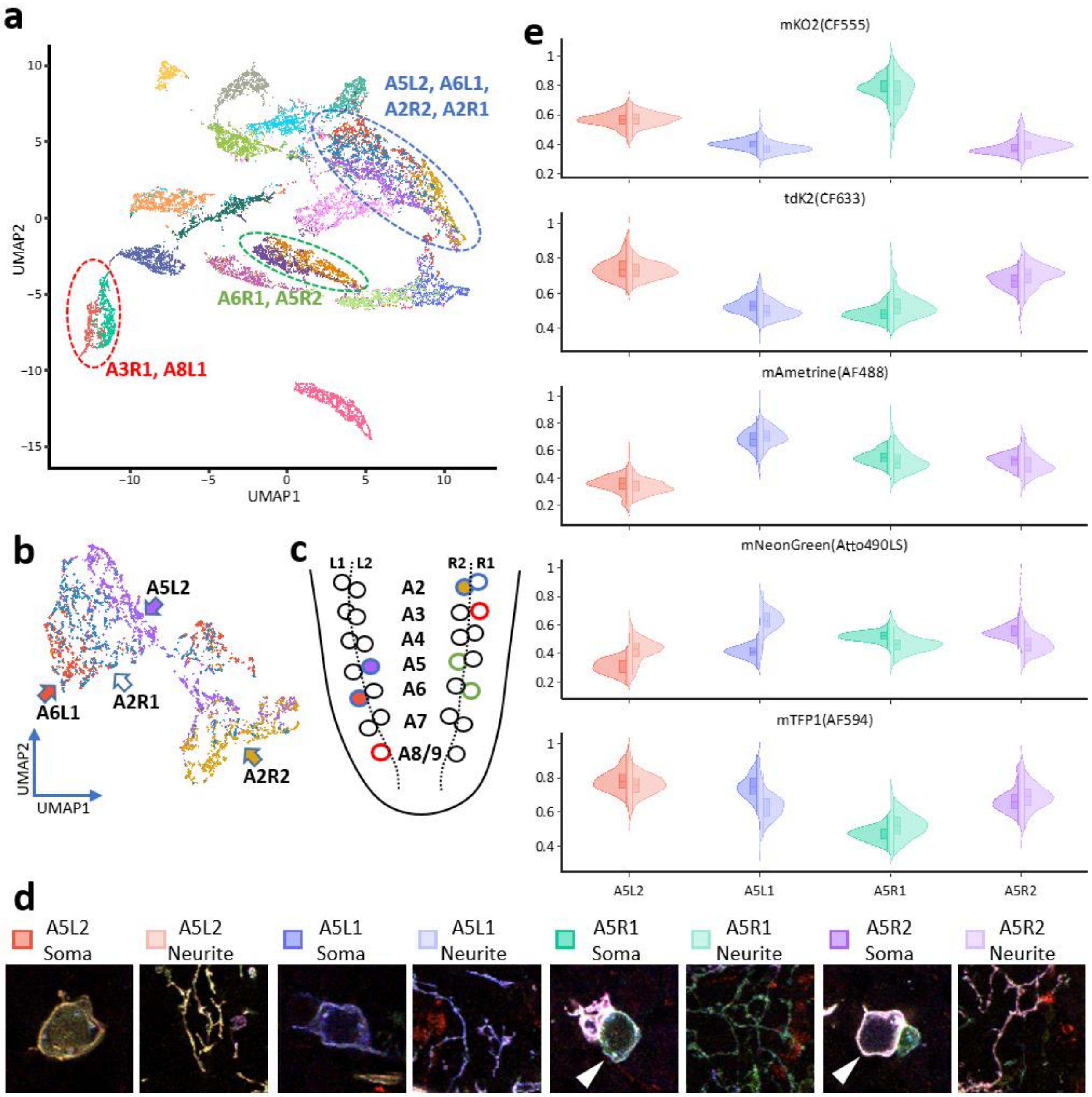
a
UMAP1
UMAP2
A5L2, A6L1,
A2R2, A2R1
A6R1, A5R2
A3R1, A8L1
b
A5L2
A6L1
A2R1
A2R2
UMAP2
UMAP1
c
L1 : L2
R2 : R1
A2
A3
A4
A5
A6
A7
A8/9
d
A5L2 Soma
A5L2 Neurite
A5L1 Soma
A5L1 Neurite
A5R1 Soma
A5R1 Neurite
A5R2 Soma
A5R2 Neurite
e
mKO2(CF555)
tdK2(CF633)
mAmetrine(AF488)
mNeonGreen(Atto490LS)
mTFP1(AF594)
A5L2
A5L1
A5R1
A5R2

**Figure S12. Bitbow color analysis of the traced 21 larval VNC serotonergic neurons. (a)** The normalized pixel intensities in each Bitbow channel along the 21 neurons' soma tracing were recorded and projected as a 2D UMAP display. 16 well-separated clusters, representing 16 resolvable Bitbow colors are identified on the UMAP embedding. 3 of the 16 clusters contain pixels sampled from more than one neuron, marked by red, green or blue dashed circles with the neuron names indicated in the same colors. Among these 3 clusters, the red- and green-circled clusters contain 2 neurons each and the two neurons composed consistent subtle spectral differences that were indicated by their tight pixel locations within the same cluster. The blue-circled cluster contains 4 neurons, whose pixel locations are much more intermingled on this UMAP. **(b)** We took those pixels belonging to the blue-circled cluster and remapped them on a higher resolution 2D UMAP display and found that 3 of the 4 neurons can be separated by consistent subtle spectral differences (solid arrows). **(c)** We marked the soma positions of the 8 neurons with less separable Bitbow colors in a schematic of the VNC serotonergic neurons. The circle outline colors correspond to the 3 dashed color line-circled clusters in (**a**). For the blue-circled neurons, their filled colors correspond to the filled colors of the arrows in (**b**). **(d)** Example images of the soma and distal neurites of the four A5 neurons displayed in **Fig. 5b**. **(e)** Half-violin plots of the four A5 neurons' normalized color intensities in each Bitbow spectral channel. Left and right halves plots pixels sampled from soma tracings and distal neurite tracings, respectively. Plotting colors correspond to the same square colors in (**d**), which indicate different A5-neurons. Boxes inside the violin plots are 1st quartile, median and 3rd quartile indicators. Y-axis, normalized fluorescence intensity in each channel.

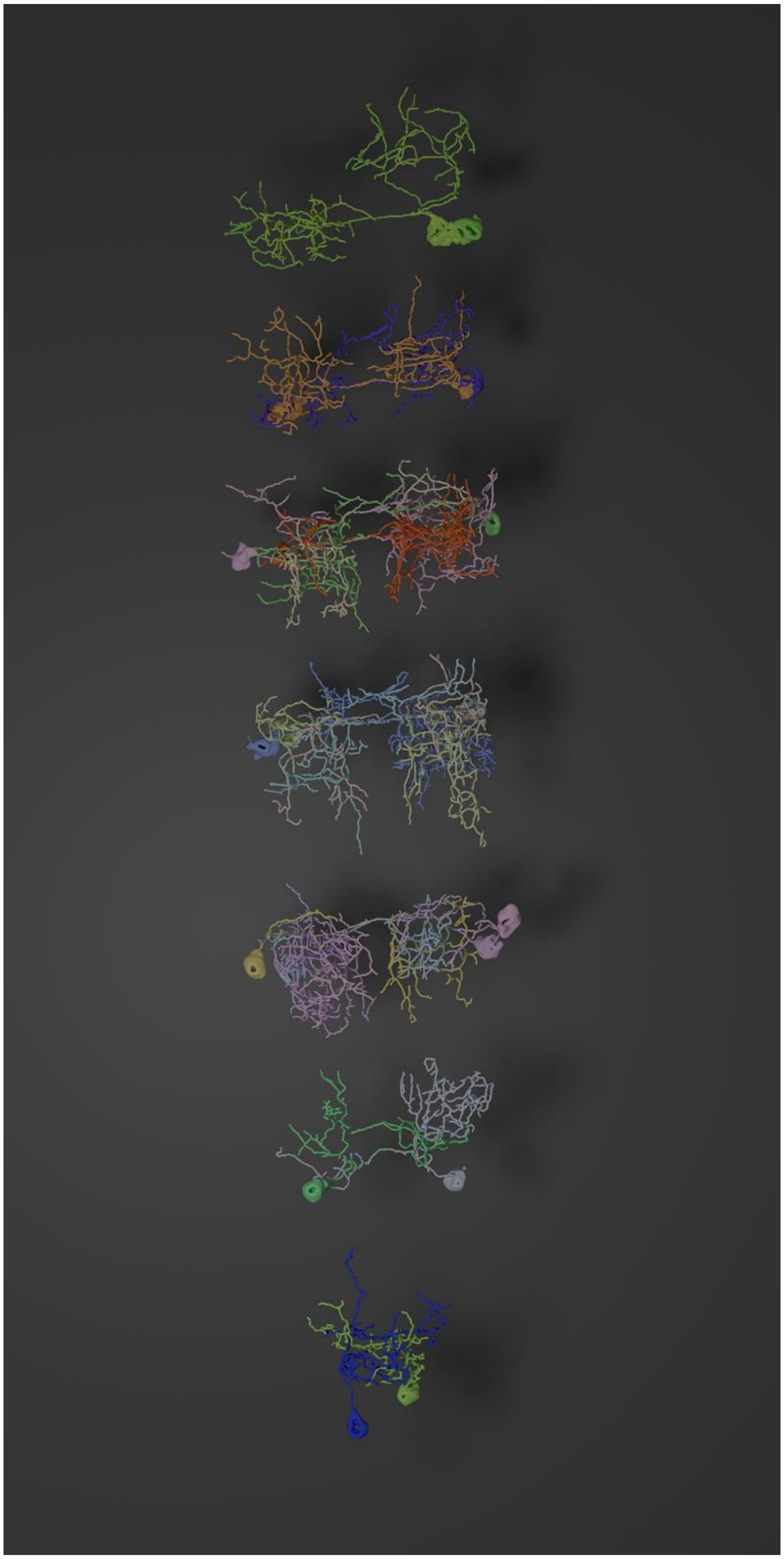

**Figure S13. Segmental display of tracings from abdominal serotonergic neurons in a *Drosophila* larvae.** Segments were organized from anterior to posterior, as the top neurons belong to segment A2, and the bottom neurons belong to segment A8.

### Movie Legends

**Movie S1. Bitbow2 labeled VNC serotonergic neurons traced by nTracer.** Three VNC hemi-segments of an offspring of the Bitbow2.1 fly and TRH-Gal4 fly cross is processed by a modified protein-retention Expansion Microscopy protocol and imaged by a confocal microscope. The nTracer results are overlaid on the fluorescence image and 3D rendering of the tracing results is shown to illustrate the relative projection patterns of the serotonergic neurons.

**Movie S2. Animations illustrating the imaging, tracing and morphology analysis of a Bitbow2 labeled fly.** First, the volumetric fluorescent data is overviewed, followed by the reveal of the nTracer-generated tracings. Finally, all neurons are then separated into manually-assigned morphological groupings, as in **Fig. 5**.
